# L-Cells are the Functional Neuropod Cell in Human Gastrointestinal Tract and are Dysregulated in Inflammatory Bowel Disease (IBD)

**DOI:** 10.1101/2025.10.30.685584

**Authors:** Ioannis Alexandros Gampierakis, Praveen Krishna Veerasubramanian, Thomas A. Wynn, Richard L. Gieseck

## Abstract

Neuropod cells are a newly discovered type of enteroendocrine cell (EEC) that connect the gut and brain functionally into one circuit. In the mouse colon, neuropod cells express various peptide hormones, such as Pyy and Glp1, presynaptic proteins, and make synaptic contacts with sensory neurons. While their function is not fully elucidated, they play a significant role in relaying signals to the brainstem upon sensing nutrients and microbial factors in the gut lumen. Their occurrence in the human gastrointestinal tract is currently not established. In this study, we showed that PYY-expressing cells (L-cells) in the human colon exhibit characteristics of neuropod cells. Utilizing advanced histological methods and confocal microscopy we found that L-cells of the healthy human colon possess distinctive morphology, express synaptic proteins, and exist proximal to sensory neurons. This agrees with our meta-analysis of single-cell RNA sequencing (scRNA-Seq) data that showed that human colonic L-cells express pre- and post-synaptic genes. As inflammatory conditions could affect colonic neuropod cells, we aimed to profile the phenotypic and transcriptional changes of neuropod cells both in human and murine colon in Inflammatory Bowel Disease (IBD) and experimental colitis, respectively. In human IBD, the abundance of neuropod cells and spatial proximity to sensory neurons were decreased in the colon of ulcerative colitis (UC) and Crohn’s disease (CD) patients. L-cells in IBD patients display genes related to innate and adaptive immunity, including antigen presentation genes suggesting a role in immune regulation. We further confirmed the effects of intestinal inflammation in neuropod cells by utilizing the DSS mouse model of colitis, where we showed that acute DSS colitis induced spatially distinct effects on the abundance of neuropod cells and impaired the synaptic connection with sensory neurons. Overall, these findings extend early murine characterizations to the human system and highlight the complex interactions between colonic neuropod cells and the enteric nervous and immune systems during inflammatory diseases.

## Introduction

Enteroendocrine cells (EECs) are a rare epithelial cell type whose primary function is to monitor the gut lumen for nutrients, microbial factors, and pathogens^1^. In response, EECs secrete a multitude of peptides, hormones, and neurotransmitters that act in autocrine and paracrine manner on neighboring cells including other EECs, epithelial cells, leukocytes, and sensory neurons^2,3^.

Recently, a new enteroendocrine cell type was identified in the mucosa of the murine small intestine and colon that exhibits both endocrine and neuronal characteristics and forms synapses via specialized extrusions with vagal nodose neurons.^4^. These extrusions with neuron-like features were named neuropods, and EECs with this morphology were named neuropod cells^4^. Further studies revealed that neuropod cells co-express various pre- and post-synaptic proteins^5^ such as synapsin-1 (Syn1) and post-synaptic density 95 (Psd95) in addition to the peptides and hormones expressed and secreted by EECs, including cholecystokinin (Cck) and polypeptide YY (Pyy). In the murine colon, neuropod cells mainly express Pyy and are increasingly recognized as a subtype of L-cells^5^, the most abundant EEC type in the murine and human colon^1^. The presence of neuropod cells in the human colon has not yet been directly investigated, but there are a few studies suggesting that human colonic L-cells express pre-and post-synaptic genes^6^ and synaptic proteins^7^.

Neuropod cells play a crucial role in sensing nutrients in the gut lumen and transmitting information to the brainstem, with recent studies highlighting their ability to differentiate and transduce stimuli from artificial sweeteners and sugars to the brain through the vagus nerve using sweet taste receptors and sodium-glucose transporters^8,9^. Additionally, neuropod cells have been suggested to regulate visceral pain via guanylyl cyclase C, an FDA-approved target for chronic constipation syndrome treatment^10^. However, their role in intestinal inflammation is less understood. A recent study has shown that small-bowel dysmotility in experimental dextran sodium sulfate (DSS) colitis can be reversed by genetically modulating EECs or administering serotonin analogs^11^. EECs are suggested to play a key role in sensing gut microbiome changes, and deficiencies in colonic EECs can alter microbiota composition, metabolism, and lead to hyperphagia and obesity^12^. Inflammatory Bowel Disease (IBD) patients exhibit lower EEC abundance^13–15^ and downregulated hormone genes such as glucagon (GCG) and PYY in the human colon^16,17^. EEC deficiency in IBD could either directly or indirectly impact immune function as it alters microbiota composition, intestinal motility and energy regulation^2^. Importantly, neuropod cells might also play a key role in relaying signals in vagal nodose neurons and in turn activate neurons in the brainstem nuclei that are known to strongly regulate immune responses^18^.

In this study, we aimed to investigate whether neuropod cells exist in the human colon. Indeed, we showed that L-cells in the human colon exhibit characteristics of neuropod cells. We found that L-cells highly express synaptophysin and are proximal to sensory neurons in the human colon. To further study neuropod cells in the human colon, we created a human colonic epithelial cell atlas where we integrated five unique single cell RNA-sequencing (scRNA-Seq) datasets and showed that L-cells are the only epithelial cell type in the human colon that highly express presynaptic genes, further complementing our histological data and confirming the presence of neuropods in the human gastrointestinal tract for the first time.

Given the absence of literature regarding the response of neuropod cells during colonic inflammation, we also profiled the cellular and transcriptional changes of neuropod cells in both the human and murine colon. In human IBD, we noted a reduction in the spatial proximity between neuropod cells and sensory neurons along with diminished cell numbers in UC and CD patients. L-cells in IBD patients upregulate genes associated with innate and adaptive immunity, including those related to antigen presentation, suggesting a potential role in immune regulation. To study the response of neuropod cells during intestinal inflammation, we utilized the DSS mouse model of colitis. We found that DSS treatment induced spatially distinct effects on abundance of neuropod cells and impaired the synaptic connection with sensory neurons in the mouse colon. Additionally, the spatial proximity of neuropod cells to T cells increased during experimental colitis. Overall, our findings highlight the complex interactions between neuropod cells and the enteric nervous and immune systems, suggesting novel roles for these cells in gut health and disease.

## Results

### L-cells in the healthy human colon exhibit characteristics of neuropod cells

L-cells are the most abundant population of EECs in the mouse and human colon, mainly expressing and secreting Pyy and Glp1^19^. In the mouse colon, neuropod cells extend cytoplasmic projections (neuropods) into the lamina propria (**Figure 1A**) and express Pyy and presynaptic proteins^5^ such as synaptophysin (Syp) and synapsin 1 (Syn1). Computational analysis of published scRNA-Seq data of isolated EECs^20^ from the stomach through the rectum of mice revealed that L-type cells express various synapse-associated genes such as *Syn1*, *Syp*, *Snap25,* and *Pclo* along the small intestine and colon (**Supplementary Figure 1A-F**). To further confirm that L-cells express synaptic proteins, we stained frozen mouse colon sections for Pyy and for common presynaptic proteins such as Syn1 and Syp (**Figure 1B and C**). We found that Pyy^+^ cells with neuropod morphology highly express Synapsin 1 (85.48±1.41%; **Figure 1D**) and Synaptophysin (97.78±1.25%, **Figure 1D**). 3D image analysis showed uniform expression of presynaptic proteins Syn1 and Syp in the cell soma and neuropod area of Pyy^+^ cells (**Figure 1E-F**). These data show that neuropod cells in the mouse colon highly express presynaptic proteins. In the human colon, L-type cells comprise less than 1% of the epithelium. Due to the sparsity and rarity of this cell type, we created a human colonic epithelial cell atlas where we integrated five published scRNA-Seq datasets^6,16,17,21,22^ (**Supplementary Figure 2A-D**). Using this atlas, we were able to show that L-type cells (**Supplementary Figure 3A and B**) highly express presynaptic genes compared to other epithelial cell types such as colonocytes and goblet cells (**Supplementary Figure 3C**). Among the highly expressed genes are the presynaptic genes *SYP*, *PCLO,* and *SNAP25* (**Supplementary Figure 3C**). To further validate the above transcriptional data, we stained healthy human colon formalin fixed paraffin embedded (FFPE) sections for PYY and SYP to probe PYY^+^ cells for presynaptic protein expression (**Figure 1G-I**). We found that 99±1.00% of PYY^+^ cells in the human colon express SYP (**Figure 1J**). Our analysis showed that most PYY^+^ cells in the human colon exhibit morphology characteristic of neuropod cells (**Figure 1K**) with extending neuropods and soma that uniformly express SYP (**Figure 1L**). These data suggest that human colonic L-cells display the characteristic neuropod morphology of mouse neuropod cells and highly express presynaptic proteins. We further noticed that subsets of human colonic L-cells display morphology not described before in mouse neuropods. More specifically, histology analysis revealed four distinct types of L-cells expressing PYY in the human colon. The first type, referred to as Type A, exhibits canonical neuropod cell morphology, with its neuropods oriented towards the lumen (**Figure 1M**). The second type, referred to as Type B, exhibits two neuropods (**Figure 1M**). One neuropod is oriented towards the lamina propria, while the other extends towards the lumen. The third type (Type C) resembles the murine neuropod cell morphology with its neuropod oriented towards the lamina propria (**Figure 1M**). The fourth type is a close L-cell type and has no neuropods (**Figure 1M**). The most abundant type of L-cells in the human colon was Type A (50.13±6.21%), followed by the close L-cell type (38.76±9.06%), Type C (11.17±6.12%), and finally Type B (6.24±4.46%) which was the rarest type (**Figure 1N**). These data suggest that human neuropod cells could be categorized into three different types and are morphologically heterogeneous with canonical neuropod polarities (**Figure 1O**). Overall, we showed for the first time that neuropod cells are present in the human colon, expressing presynaptic genes and proteins with canonical neuropod polarities projecting either in the lumen or the lamina propria.

**Figure 1.**
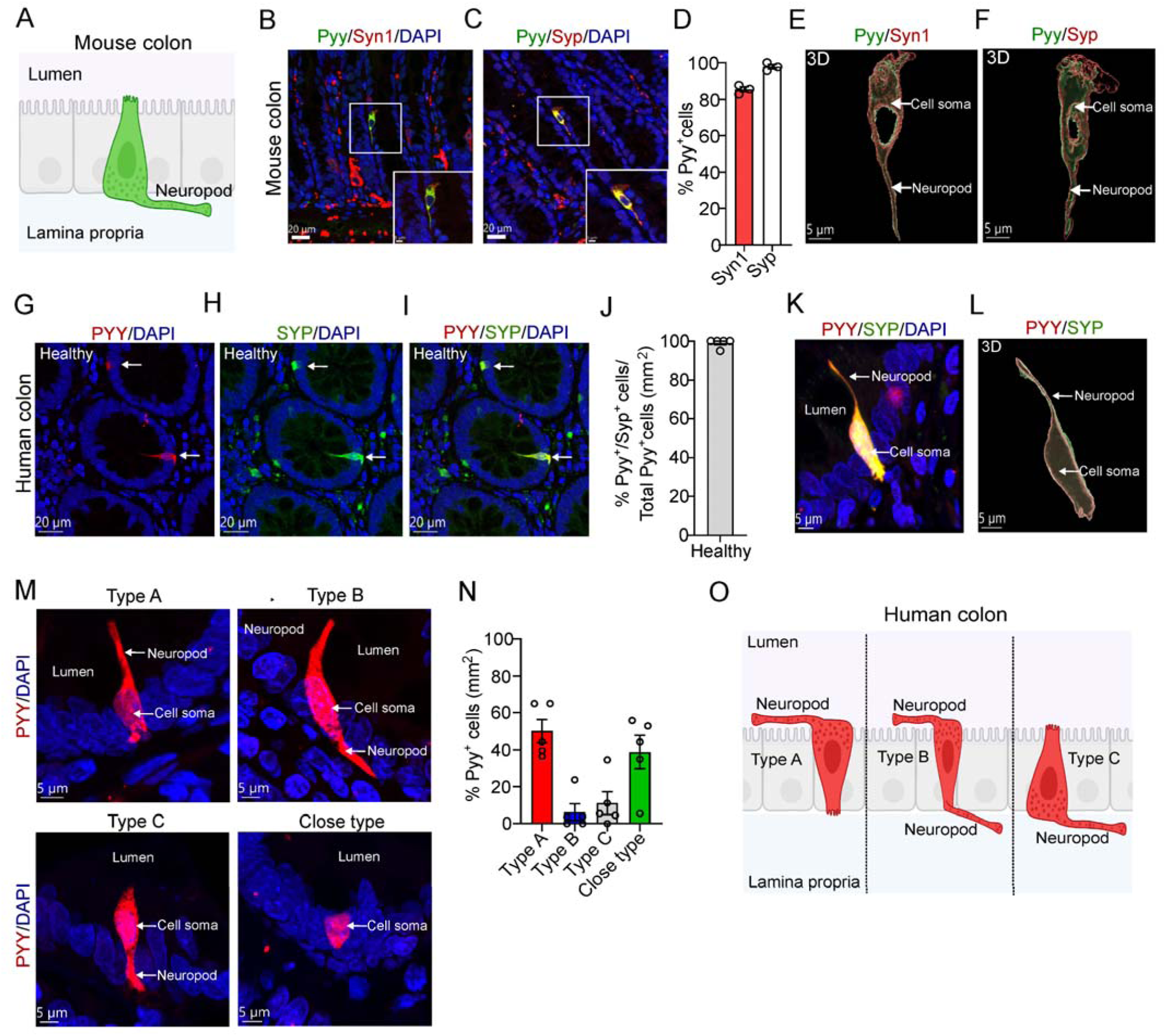
Human colonic L-cells display neuropod-like characteristics. (A) Schematic of a neuropod cell in the mouse colon. (B-C) Representative confocal images of 10 μm thick frozen mouse colon section stained for Pyy Syn1 and Syp. DAPI, 4,6-diamidino-2-phenylindole. Scale bars, 20 μm. Insets: Magnified confocal images of Pyy^+^ cells expressing Syn1 and Syp respectively. (D) Percentage of Pyy^+^ cells with neuropod morphology expressing Syn1 and Syp (n=3 mice). (E-F) 3D image reconstruction of Pyy^+^ cells with neuropod morphology showing the even distribution of Syn1 and Syp both in the cell soma and neuropod area. Scale bars, 5 μm (G-I) Representative confocal images of 5 μm FFPE human colon sections stained for PYY and SYP. DAPI, Scale bar 20 μm (J) Percentage of PYY^+^ cells expressing SYP normalized to the total number of PYY^+^ cells per mm^2^ (n=5 human colon samples). (K-L) Representative magnified confocal image and 3D reconstructed image of individual PYY^+^ cells stained for SYP, depicting even distribution of PYY and SYP in the cell soma and neuropod area. DAPI, Scale bar 5 μm (M) Representative confocal images of 5 μm thick FFPE human colon sections stained for PYY depicting three types of neuropod polarities named as Type A, Type B and Type C. Close type PYY-expressing cells do not extend neuropods. DAPI, Scale bar 5 μm (N) Percentage of the various types of PYY^+^ cells normalized to the total number of PYY^+^ cells per mm^2^ (n=5 human colon samples). (O) Schematic of the different types (Type A, B and C) of neuropod polarities in the human colon. Plotted are means± SEM. Schematics were created in BioRender (https://BioRender.com/x7tmjdx)

### L-cells in the healthy mouse and human colon are proximal to sensory nerve terminals

As previously described, neuropod cells form synaptic connections with sensory neurons in the mouse colon^5^. To assess whether neuropod cells and L-cells synapse with sensory neurons in the mouse colon, we stained the whole mouse colon for Pyy and for the common marker of sensory neurons Pgp9.5. We observed that most Pyy^+^ cells were proximal to Pgp9.5^+^ sensory neurons independent of the presence or absence of neuropods (**Figure 2A and B**). We then quantified the distance between Pyy^+^ cells and Pgp9.5^+^ sensory neurons by 3D image analysis (**Figure 2C-E**). We found that across all regions of the mouse colon (proximal, mid and distal) the relative frequency of Pyy^+^ cells in proximity to Pgp9.5^+^ sensory neurons (Distance 0 μm) was about 77% (relative frequency range from 71 to 82%) (**Figure 2F-H).** 3D image analysis also showed that the sensory nerve terminals were proximal to the soma area of the cell (**Figure 2D and Supplementary Figure 4A and B**) in both Pyy^+^ cells irrespective of the presence of neuropods (**Supplementary Figure 4A and B**). The above data suggests that the vast majority of neuropod cells are proximal to sensory neurons in the mouse colon. To assess whether human L-cells and neuropod cells are proximal to sensory neurons, we stained human colon FFPE sections for PYY and PGP9.5 (**Figure 2I**). 3D image analysis revealed that the sensory nerve terminals are proximal to the soma area of neuropod cells (**Figure 2J**) and nearly 50% of the PYY^+^ cells were in proximity (Distance 0 μm) to PGP9.5^+^ sensory neurons (**Figure 2K and L**). These data suggest that human colonic neuropod cells are proximal to sensory nerve endings suggesting a novel neuroepithelial network.

**Figure 2.**
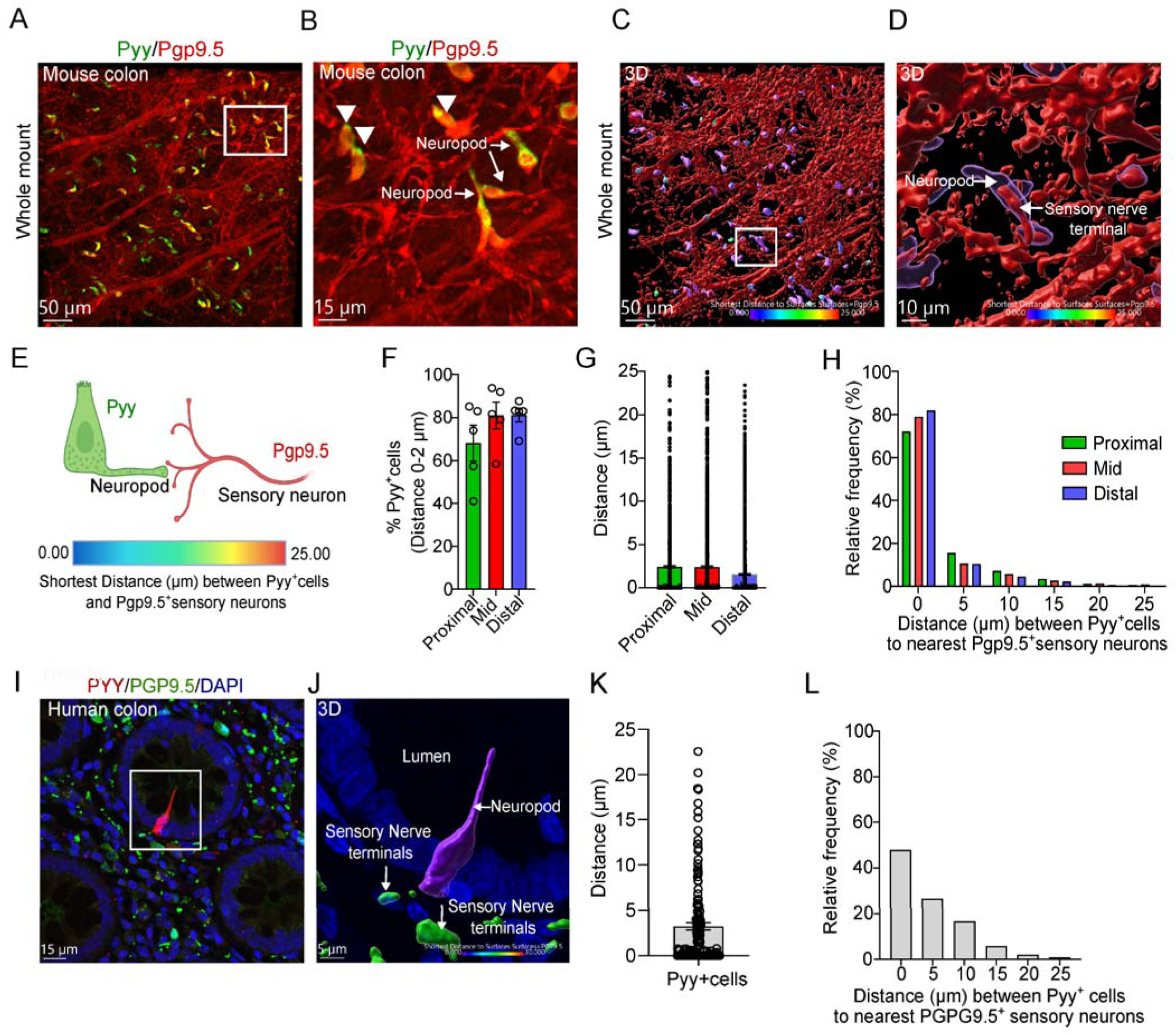
Human and mouse colonic neuropod cells are proximal to sensory neurons. (A-B) Representative confocal image of whole mount colon stained for Pyy and Pgp9.5. Scale bar 50 μm. Inset: Magnified confocal image depicting proximity of Pyy^+^ cells with neuropod and non-neuropod morphology to Pgp9.5^+^ sensory neurons. Scale bar 15 μm (C-D) Representative 3D image of whole mount colon stained for Pyy and Pgp9.5. Scale bar 50 μm. Inset: Magnified 3D image depicting proximity between Pyy^+^ cells with neuropod morphology and Pgp9.5^+^sensory neurons. Scale bar 10 μm (E) Schematic depicting 3D imaging process for quantifying the shortest distance between Pyy^+^ cells and Pgp9.5^+^sensory neurons; purple to blue colon indicates distance less than 2 um, green to red indicates distance close to 20-25 um. Created in BioRender (https://BioRender.com/p0s59mm) (F) Percentage of Pyy^+^ cells in proximity (Distance 0 μm) to Pgp9.5^+^ sensory neurons across colon regions (proximal, mid and distal). n=5-6 mice (G) Graph illustrates the distance between Pyy^+^ cells (pooled from 5 mice) and the nearest Pgp9.5^+^sensory neurons (H) Relative frequency of distance (μm) between Pyy^+^ cells and nearest Pgp9.5^+^sensory neurons across colon regions, pooled data from 5 mice. (I) Representative confocal image of 5 μm thick FFPE human colon section stained for PYY and PGP9.5. DAPI, Scale bar 15 μm (J) Magnified 3D image showing the distance between PYY^+^ cells and PGP9.5^+^sensory neurons. DAPI, Scale bar 5 μm (K) Graph illustrates the distance between PYY^+^ cells (pooled from 5 human colon samples) and the nearest PGP9.5^+^sensory neurons. (L) Relative frequency of distance (μm) between PΥΥ^+^ cells and nearest PGP9.5^+^sensory neurons, pooled data from 5 human colon samples. Plotted are means± SEM.

### Experimental DSS colitis and IBD decrease the density of L-cells in the human and mouse distal colon respectively

Colonic EECs are crucial for sensing gut microbiota and their metabolites, secreting peptide hormones and cytokines in response^2^. Dysbiosis is known to occur in IBD along with an excessive inflammatory response and destruction of the epithelial lining^23^. There is an increasing consensus suggesting that colonic EECs might play a significant role in immune regulation and epithelial homeostasis^2^.

In light of this, we first examined whether the density of mouse neuropod cells is affected by experimental colitis. To do so, we utilized the dextran sodium sulfate (DSS) murine model of colitis^24^. Acute colitis was induced by administering 3.5% w/v DSS in the drinking water for seven consecutive days (**Figure 3A**). The mice were sacrificed three days later, at the ten-day time point. Mice treated with DSS experienced significant weight loss (**Figure 3B**), severe intestinal inflammation evident by increased fecal levels of lipocalin 2 (**Figure 3C**), and decreased colon length (**Figure 3D**). DSS treatment in mice induces a robust inflammatory response in the distal colon^25^. Notably, we found an increase in the protein levels of proinflammatory cytokines such as Il1b, Il11, Il6, Tnfa, Csf1, and Osm (**Supplementary Figure 5A-C**), chemokines such as Ccl2 and Ccl3 (**Supplementary Figure 5A-C**), pro-inflammatory adipokines such as Lcn2, Rbp4, Ob, and Adsf (**Supplementary Figure 5A-C**), and pentraxins such as Crp, Ptx2, and Ptx3 (**Supplementary Figure 5A-C**) in the distal colon of DSS-treated mice. Similarly, we found an increase in various proinflammatory cytokines, chemokines, proinflammatory adipokines and pentraxins in the proximal and mid colon of DSS treated mice (**Supplementary Figure 5 A-C**).

**Figure 3.**
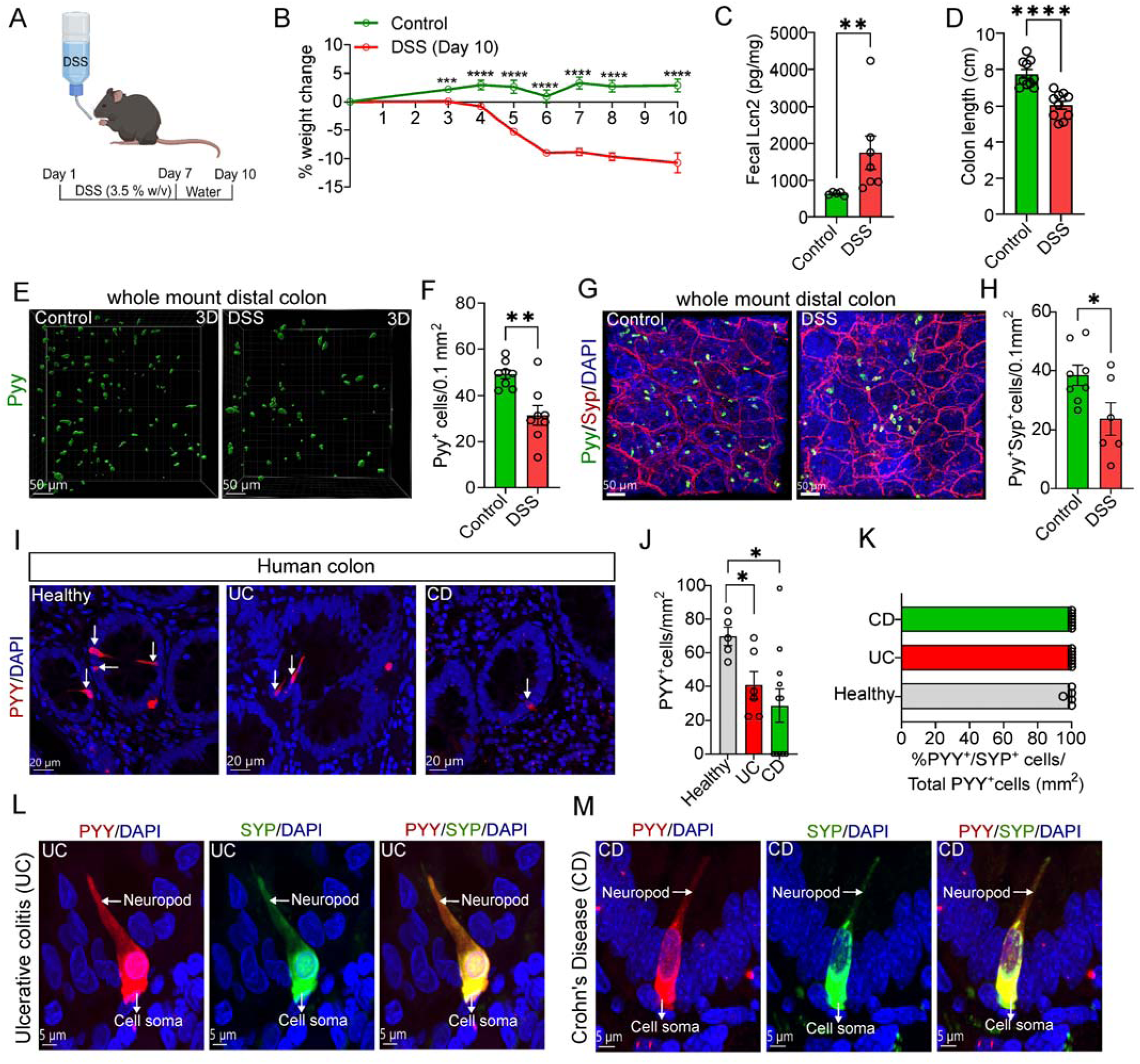
Acute DSS colitis and IBD decreased neuropod cells’ abundance in the mouse distal colon and human colon respectively. (A) Schematic depicting the mouse model of DSS-induced colitis. Mice received 3.5% w/v DSS in their drinking water for 7 days and sacrificed 3 days later, marking a 10-day time period. Created in BioRender (https://BioRender.com/p68n398) (B-D) Colitis readouts, including weight loss; fecal lipocalin-2 (Lcn2) levels, and colon length measurements (n= 5-10 mice/group). (E) Representative 3D images of whole mount distal colon stained for Pyy of control and DSS-treated mice. Scale bar 50 μm (F) Quantification of Pyy^+^ cells per 0.1 mm^2^ in the distal colon of control and DSS-treated mice (n= 8 mice/group). (G) Representative confocal images of whole mount distal colon stained for Pyy and Syp in control and DSS-treated mice. DAPI, Scale bars, 50 μm (H) Quantification of Pyy^+^ Syp^+^ cells per 0.1 mm^2^ in the distal colon of control and DSS-treated mice (n=6-8 mice/group). (I) Representative confocal images of 5 μm thick FFPE human colon sections of healthy subjects, UC and CD patients stained for PYY. DAPI, Scale bar 20 μm (J) Quantification of PYY^+^ cells per mm^2^ in FFPE human colon sections of healthy subjects (n= 5), UC patients (n=6), and CD patients (n=11). (K) Percentage of PYY^+^ cells expressing SYP normalized to the total number of PYY^+^ cells per mm^2^ in the colon of healthy subjects (n=5), UC patients (n=6), and CD patients (n=6). (L-M) Representative magnified confocal images of 5 μm thick FFPE human colon section of UC and CD patients stained for PYY and SYP, depicting even distribution of PYY and SYP in the cell soma and neuropod area. DAPI, Scale bar 5 μm Plotted are means± SEM. *p<0.05, **p<0.01, ***p<0.001, ****p<0.000 by unpaired two-tailed tests and body development over time was evaluated by multiple unpaired two-tailed t-tests (one per timepoint) with Holm-Sidak correction. The significance of the differences between more than two groups was evaluated using ANOVA followed by Kruskal-Walli’s comparison test. Experiments were repeated at least twice.

As distal colon was the most affected part of the colon in DSS-treated mice, we performed immunofluorescence for Pyy in whole-mount distal colon preparations from both control and DSS-treated mice. Our 3D image analysis (**Figure 3E**) showed decreased density of Pyy^+^ cells in the distal colon of DSS-treated mice (**Figure 3F**). Moreover, when we evaluated the impact of DSS treatment on the expression of presynaptic proteins in neuropod cells in the distal colon, we found decreased density of Pyy^+^/Syp^+^ cells in the distal colon of DSS-treated mice (**Figure 3G and H**). However, when we examined other colon parts such as the proximal and mid colon, we found an unexpected increase in the density of Pyy^+^ in the proximal colon of DSS-treated mice (**Supplementary Figure 5D and E**), while no effects were observed in the density of Pyy^+^ cells in the mid colon of DSS-treated mice (**Supplementary Figure 5F and G**). Similarly, the density of Pyy^+^/Syp^+^ cells was increased in the proximal colon of DSS-treated mice (**Supplementary Figure 5H and I**), while no differences in the density of Pyy^+^/Syp^+^ cells were observed in the mid colon of DSS-treated mice (**Supplementary Figure 5I**). These data suggest that DSS treatment induces spatially distinct effects on neuropods cells’ abundance along the mouse colon epithelium.

To evaluate the translational implications of the aforementioned findings, we investigated whether the density and phenotype of human colonic neuropod cells are affected in patients with IBD. Histology analysis of FFPE colon samples from IBD patients (including UC and CD patients) showed decreased density of PYY^+^ cells in the colon of UC and CD patients compared to healthy subjects (**Figure 3I and J**). However, no differences were found in the percentages of Type A (**Supplementary Figure 6A**), Type B (**Supplementary Figure 6B**), Type C (**Supplementary Figure 6C**), and close type L-cells (**Supplementary Figure 6D**) between healthy, UC and CD patients. Furthermore, we explored the potential changes in the expression of presynaptic proteins, such as SYP in neuropod cells in the colon of IBD patients. Staining for PYY and SYP showed no changes in the expression of SYP in PYY^+^ cells in the colon of UC and CD patients when compared to healthy individuals (**Figure 6K**). Interestingly, the colocalization analysis in the colon sections from UC and CD patients showed that 100% of the PYY^+^ cells express SYP (**Figure 6L and M**). Transcriptionally, there was no significant difference in the expression of more than 200 pre- and postsynaptic genes in L-cells when comparing healthy with involved and non-involved IBD samples. (**Supplementary Figure 6E**). These data suggest that the density of neuropod cells is decreased in IBD, while the expression of synaptic genes and proteins remains unaffected.

### Experimental DSS colitis and IBD affect the proximity of neuropod cells to sensory neurons along the colon epithelium

To further identify changes in neuropod cells’ phenotypes during experimental DSS colitis and IBD, we evaluated the spatial changes in the proximity of neuropod cells to sensory neurons. For that purpose, we stained whole mount preparations of the proximal, mid, and distal regions of the colon for Pyy and Pgp9.5 from both the control and DSS-treated mice. 3D image analysis (**Figure 4A and B**) showed that DSS treatment decreased the relative frequency of Pyy^+^ cells in proximity to Pgp9.5^+^ sensory neurons in the mid colon by 26% (**Figure 4C and D**). Similarly, we found a trending decrease in the relative frequency of Pyy^+^ cells in proximity to Pgp9.5^+^ sensory neurons in the distal colon, but it was not statistically significant (**Supplementary Figure7 A**). There were no changes in the relative frequency of Pyy^+^ cells in proximity to Pgp9.5^+^ sensory neurons in the proximal colon between the control and DSS-treated mice (**Supplementary Figure 7B**). These data suggest that acute DSS colitis induces distinct effects on the spatial proximity between neuropod cells and Pgp9.5^+^ sensory neurons, mainly affecting the mid and distal colon in DSS-treated mice.

**Figure 4.**
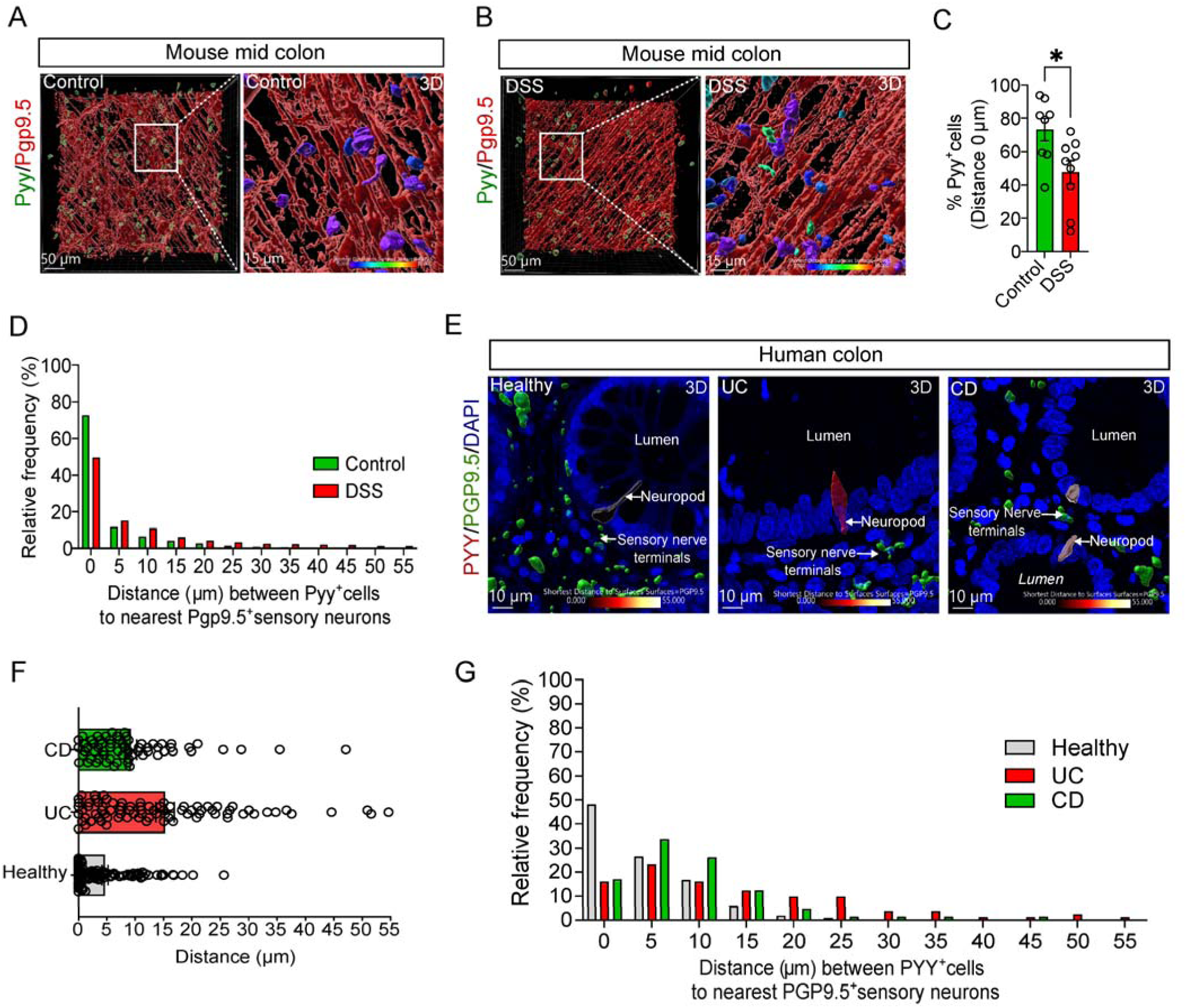
Acute DSS colitis and IBD altered the proximity of neuropod cells to sensory neurons. (A-B) Representative 3D image of whole mount mid colon from control and DSS-treated mice stained for Pyy and Pgp9.5. Scale bar 50 μm. Inset: Magnified 3D image depicting proximity between Pyy^+^ cells and Pgp9.5^+^sensory neurons. Scale bar 15 μm (C) Percentage of Pyy^+^ cells in proximity (Distance 0 μm) to Pgp9.5^+^ sensory neurons in the mid colon of control (n= 8) and DSS-treated (n=9) mice. (D) Relative frequency of distance (μm) between Pyy^+^ cells and nearest Pgp9.5^+^sensory neurons in the mid colon of control (pooled data from 8 mice) and DSS-treated mice (pooled data from 9 mice). (E) Representative 3D images depicting the distance between PYY^+^ cells and PGP9.5^+^sensory nerve terminals in healthy subjects, UC and CD patients. DAPI, Scale bar 10 μm (F) Graph illustrates the distance between PYY^+^ cells and the nearest PGP9.5^+^sensory neurons in healthy subjects (pooled data from 5 samples), UC patients (pooled data from 6 samples), and CD patients (pooled data from 6 samples). (G) Relative frequency of distance (μm) between PΥΥ^+^ cells and nearest PGP9.5^+^sensory neurons in healthy subjects (pooled data from 5 samples), UC patients (pooled data from 6 samples), and CD patients (pooled data from 6 samples). Plotted are means± SEM. *p<0.05 by unpaired two-tailed tests. Experiments were repeated at least twice.

To evaluate the impact of human IBD on the synaptic connectivity of neuropod cells with sensory neurons, we performed staining for PYY and PGP9.5 in FFPE colon sections of healthy and IBD patients. 3D image analysis (**Figure 4E**) revealed a 40% decrease in the relative frequency of PYY^+^ cells in proximity to PGP9.5^+^sensory neurons (Distance 0 um) in the colon of IBD patients (both UC and CD) compared to heathy individuals (**Figure 4F and G**). These data suggest that IBD decreases the synaptic connection between neuropod cells and sensory neurons in the human colon.

### Increased expression of antigen presenting genes in neuropod cells in the colon of IBD patients

Using the human colonic epithelial cell atlas, we were able to show that neuropod cells upregulate many genes associated with innate and adaptive immunity in IBD (**Figure 5A and B**). Among the upregulated genes were genes related to antigen presentation such as *HLA-DPA1*, *HLA-DRB1*, *HLA-DRA*, and *HLA-DPB1* (**Figure 5A; Supplementary Figure 8A**). Additionally, we found an increased gene expression of *LCN2*, *LYZ,* and *IGHA1*, genes associated with antimicrobial activity and immune defense (**Figure 5A and B**). These data suggest that colonic neuropod cells might play a role in immune regulation in IBD.

**Figure 5.**
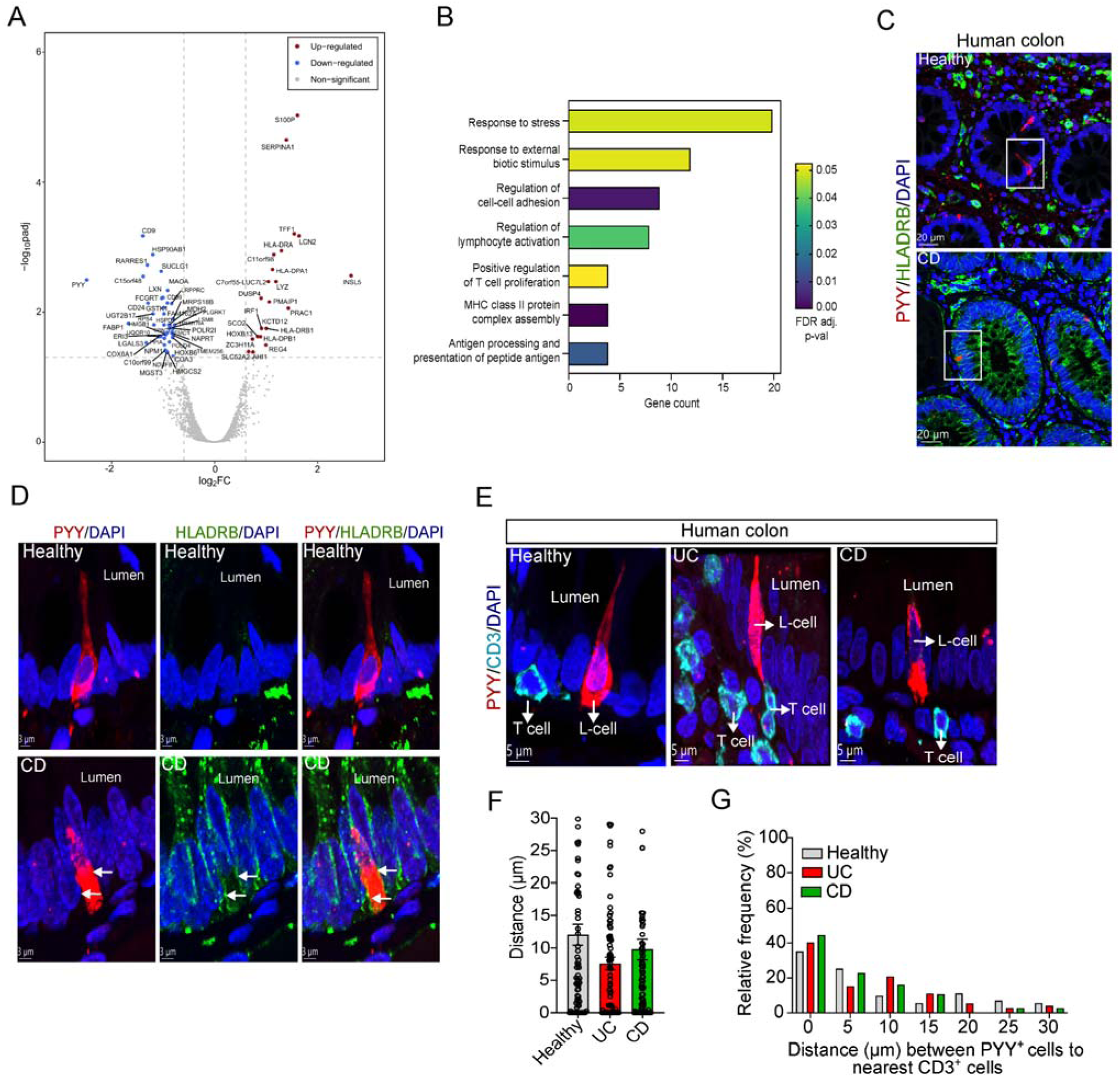
Human colonic L-cells upregulate genes related to antigen presentation in IBD. (A) Volcano plot of differentially expressed genes in L-cells in colon of healthy vs inflamed colon of IBD patients. (B) Gene ontology overrepresentation analysis of the genes differentially expressed in L-cells from IBD samples. (C-D) Representative confocal image of 5 μm thick FFPE human colon section from a healthy subject and a CD patient stained for PYY and HLA-DRB. DAPI, Scale bar 20 μm. Inset: magnified confocal images stained for PYY and HLA-DRB. White arrows depict HLA-DRB expression in the cell membrane of the neuropod cell. DAPI, Scale bar 3 μm (E) Magnified representative confocal images of 5 μm thick FFPE human colon sections from healthy subjects, UC, and CD patients stained for PYY and CD3-marker for T cells. Scale bar 5 μm (F) Graph illustrates the distance between PYY^+^ cells and the nearest CD3^+^ T cells in healthy subjects (pooled data from 5 samples), UC patients (pooled data from 6 samples), and CD patients (pooled data from 6 samples). (G) Relative frequency of distance (μm) between PΥΥ^+^cells and nearest CD3^+^ T cells in healthy subjects (pooled data from 5 samples), UC patients (pooled data from 6 samples), and CD patients (pooled data from 6 samples). Plotted are means± SEM.

Among the gene ontology terms that are associated with the genes differentially expressed in L-type cells in IBD are “antigen processing and presentation of peptide antigen”, “MHC class II protein complex assembly”, “positive regulation of T cell proliferation”, and “Regulation of lymphocyte activation” **(Figure 5B)**. Prior studies have shown histologically and transcriptomically that epithelial cells in active IBD patients exhibit upregulated expression of HLA molecules^17,26^. Therefore, we asked whether human colonic neuropod cells could play a role in antigen presentation by directly interacting with T cells. The first step was to see if neuropod cells express antigen presentation proteins in their cell membrane. We stained human colon sections from healthy subjects and IBD patients for PYY, HLA-DRB, and HLA-DRA which are common markers for antigen presenting cells. As expected, we found most HLA-DRB^+^ cells in the lamina propria of healthy subjects and no expression of HLA-DRB in PYY^+^ cells (**Figure 5C and D**). In contrast, the expression of HLA-DRB was increased in the epithelium and lamina propria of IBD patients (**Figure 5C and D**), while some but not all PYY^+^ cells expressed HLA-DRB in the colon of IBD patients (**Figure 5D; Supplementary Figure 8B-E**). Interestingly, we observed HLA-DRA expression in some but not all PYY^+^ cells in healthy colon sections (**Supplementary Figure 8F-H**), while in IBD colon sections, HLA-DRA expression was not evident in PYY^+^ cells (**Supplementary Figure 8I-N)**. These differences suggest that the expression of antigen presenting proteins in neuropod cells is differentially regulated in health and IBD.

In IBD, the proximity between epithelial cells and T cells is crucial in driving chronic inflammation^27,28^. A compromised epithelial barrier allows antigens and pathogens to penetrate, activating T cells in the lamina propria. Chronic activation of mucosal T cells could lead to the disruption of tolerance mechanisms^29^, leading to excessive immune activity and the secretion of proinflammatory cytokines leading to further damage of epithelial cells^30^. Given the role of L-cells in sensing bacteria and various antigens^12^, and the fact that L-cells express antigen-presenting proteins in health and IBD, we explored whether neuropod cells closely interact with T cells in IBD. To achieve this, we stained human colon sections from healthy and IBD patients for Pyy and CD3, a marker of T cells. Then, we performed 3D imaging analysis and measured the proximity of PYY^+^ cells to CD3^+^ cells (**Figure 5E**). Interestingly, we found that in the colon of IBD patients, the proximity of PYY^+^ cells to CD3^+^ cells was not increased, compared to healthy subjects (**Figure 5F**). Despite that, we found that even in healthy colon 30% of PYY^+^ cells were proximal to CD3^+^ cells (**Figure 5G**). Overall, these data suggest that human colonic neuropod cells express antigen presenting proteins in their cell membrane and closely interact with T cells in health and IBD.

To validate the above observations in the DSS mouse model of colitis, we first checked whether L-cells express antigen presentation proteins in DSS-treated mice. For that purpose, we stained whole mount colon of control and DSS-treated mice for Pyy and MHCII. Interestingly, colocalization analysis did not show any expression of MHCII in Pyy^+^ cells. MHCII expression was primarily detected in other epithelial cell types and cells in the lamina propria in DSS-treated mice (**Supplementary Figure 9A and B**). To see whether neuropod cells closely interact with T cells in control and DSS-treated mice, we stained whole mount preparations of the proximal, mid, and distal colon for Pyy and Cd3 (**Supplementary Figure 9C**). Interestingly, we found a consistent increase (up to 50%) in the proximity of Pyy^+^ cells to Cd3^+^ cells in the proximal, mid, and distal region of the colon of DSS-treated mice (**Supplementary Figure 9D and E**). In addition, the proximity of L-cells to T cells in control mice was between 10 to 30% (Distance 0 µm), depending on the region of the colon (**Supplementary Figure 9F-H)**. These data suggest that acute DSS colitis does not induce the expression of antigen presentation proteins in L-cells but increases their spatial proximity to T cells in the different regions of the colon.

## Discussion

In this study, we present for the first time evidence of the presence of neuropod cells in the healthy human colon and evaluated their response in IBD and experimental colitis. Here, we showed that L-cells in the human colon exhibit characteristics of neuropod cells. More specifically, we showed that the majority of colonic L-cells highly express the synaptic protein synaptophysin and are proximal to sensory neurons. Moreover, using the human colonic epithelial cell atlas, we showed that colonic L-cells highly express pre-and postsynaptic genes in the healthy human colon compared to other epithelial cell types such as colonocytes and goblet cells. These data agree with our mouse data that show high expression of the synaptic proteins Synapsin 1 and Synaptophysin in L-cells of the mouse colon and close spatial proximity to sensory neurons in the different regions of the mouse colon. These data suggest that human neuropod cells exhibit the cellular and phenotypic characteristics of murine neuropod cells.

One of the main differences we observed in human neuropod cells compared to mouse neuropods is that they exhibit different neuropod morphologies. The main morphological characteristic of murine neuropod cells is the extension of neuropods towards the lamina propria. Here, we identified three types of neuropod polarities and named them as Type A, Type B and Type C. The most abundant type of neuropod cells was Type A, with a neuropod oriented towards the lumen, while only Type C resembles the polarity of murine neuropods and Type B exhibits two neuropods, one projecting towards the lumen and the other towards the lamina propria. These data suggest the morphological heterogeneity of neuropods cells in the human colon, reflecting the complexity of neuropods cell’s potential role in intestinal homeostasis.

Colonic EECs are suggested to play a role in microbial sensing, energy regulation, colon motility, and visceral pain^1^. Specifically, GUCY2C-expressing neuropod cells control visceral hypersensitivity and might play a key role in pain control in Irritable Bowel Syndrome (IBS)^10^. However, the role of neuropod cells in intestinal inflammation is poorly understood. Here, we evaluated the response of neuropod cells both in experimental colitis and IBD. Using the DSS mouse model of colitis we found that the abundance of neuropod cells (Pyy^+^/Syp^+^) decreased in the distal colon of DSS-treated mice. A recent study has shown that DSS treatment decreases the number of EECs in the small intestine and causes small-bowel hypomotility^11^. Using transgenic mouse models to manipulate the density of EECs, the authors showed that small bowel hypomotility was associated with the reduced density of EECs^11^. Notably, mice lacking EECs showed increased small-bowel hypomotility when exposed to DSS, while mice overexpressing EECs showed improved small intestinal motility^11^. The decrease in the density of neuropod cells seen in the distal colon of DSS-treated mice could suggest a compensatory mechanism to regulate colon motility in response to increased inflammation. Additional studies in this area might be of great interest to elucidate the potential role of neuropod cells in colon-motility.

In agreement with the mouse data, the abundance of neuropod cells (PYY^+^) was decreased both in patients with UC and CD. Our data confirms previous reports of reduced PYY levels in the mucosa of UC and CD patients^13,15,31^. Interestingly, the percentage of the various types of neuropod cells (Type A, B and C) in IBD did not change significantly compared to healthy subjects, suggesting that IBD did not affect or alter the polarity of the three types of neuropod cells in the human colon. We also showed that the expression of the presynaptic protein SYP in PYY^+^ cells remained unchanged in the colon of UC and CD patients, suggesting that IBD did not affect the functional components of the synaptic machinery of neuropod cells. This finding was backed up by our transcriptional analysis using the human colonic epithelial cell atlas of more than two hundred pre-and postsynaptic genes, where we found no changes in their gene expression in the colon of healthy, involved and non-involved IBD samples.

The most striking impact of IBD on neuropod cells was that it altered their proximity to Pgp9.5^+^ sensory neurons. More specifically, we found less neuropod cells proximal to sensory neurons in the colon of both UC and CD patients compared to healthy subjects. This data suggest that inflammation could disrupt the functional synaptic connectivity between neuropod cells and sensory neurons. Similarly, we found that acute DSS colitis decreased the spatial proximity of neuropod cells and Pgp9.5^+^sensory neurons mainly in the mid colon and slightly in the distal colon. These data suggest that experimental colitis affects the synaptic connectivity between neuropod cells and sensory neurons. Interestingly, chronic DSS treatment has been shown to induce colonic immobility, especially in its middle part despite stronger contractions seen in proximal and distal ends^32^. Recent studies have shown that bacterial products such as short-chain fatty acids could stimulate GLP-1 secretion from L-cells, resulting in the acceleration of colonic peristalsis^33^. Given that neuropod cells are exposed to bacterial products and pathogens during DSS colitis, altered synaptic connectivity with sensory neurons could potentially contribute to differences in colonic immobility observed in DSS-treated mice.

Patients with metabolic and gastrointestinal diseases often present with microbial dysbiosis^34^ which can initiate cellular and transcriptional changes in EECs. Dysbiosis has been implicated in IBD^23,35^, known to significantly impact epithelial homeostasis and immune regulation. EECs could be considered the first line of “immune defense”, as they sense pathogens and microbial factors and through their secretome can initiate an immune response and activate neighboring T cells and macrophages^2^. Our differential gene expression analysis revealed that L-cells in involved samples from IBD patients highly upregulate genes related to antigen presentation (e.g. *HLA-DPA1*, *HLA-DRB1*, *HLA-DRA*, and *HLA-DPB1*). As we have already discussed above, mucosal epithelial cells could act as non-conventional presenting cells in the inflamed gut^36^. Here, we showed that a fraction of neuropod cells expresses HLA-DRB in the colon of IBD patients, while HLA-DRA is expressed only in a fraction of neuropod cells in the healthy colon, suggesting the complex and heterogeneous role of neuropod cells in immune regulation both in health and IBD. This was captured by the fact that the proximity of CD3^+^ T cells to PYY^+^ cells was relatively high in the healthy colon, suggesting close and active interaction between T cells and neuropod cells. In contrast, no changes were observed in the proximity of CD3^+^ T cells to PYY^+^ cells in the colon of IBD patients. In the interpretation of our findings, we should consider that most IBD colon samples were acquired by patients that were already under treatment (biologics or corticosteroids) that is known to halt most of the innate and adaptive immune system’s responses. Therefore, these findings cannot sufficiently determine that neuropod cells could act as non-professional antigen presenting cells both in health and IBD. Further studies using organoids and co-cultures with T cells could determine whether neuropod cells can act as non-conventional presenting cells in various inflammatory conditions. Additionally, in the DSS mouse model of colitis, the proximity of CD3^+^ T cells to neuropod cells was increased in all parts of the murine colon, suggesting that T cells closely interact with neuropod cells in the mouse colon during DSS colitis. Despite that, neuropod cells did not express the MHC class II molecules either in control or DSS-treated mice, suggesting that the close interaction of T cells and neuropod cells involves other mechanisms that need to be uncovered in future studies.

Overall, this study highlights for the first time the presence of neuropod cells in the human colon. Neuropod cells dynamically adapt during DSS colitis and IBD and interact with neighboring sensory neurons and T cells. Even though neuropod cells express several immune-related genes, their role in immune regulation in health and IBD seems complex and needs further investigation. Moreover, we showed the dynamic modification of the expression of PYY in IBD and DSS colitis, that could play a key role in colon motility and microbiome regulation. The study opens new avenues for further research into novel cellular targets for modulating intestinal inflammation in IBD.

## Methods

### Mice

8-10-week-old male C57Bl/6J mice (stock no.000664) were purchased from the Jackson Laboratory and allowed to acclimatize to the animal facility environment for at least 2 weeks before being used for experimentation. Mice were maintained on Inotiv 2916 rodent diet (Teklad global 16% protein) and Innovive chlorinated water (M-WB-300C) and housed in a light (12 h light/12 h dark) and temperature (25° C) controlled environment. All procedures were carried out and approved with the Pfizer Institutional Animal Care and Use Committee (IACUC) regulations and established guidelines.

### Human colon tissue samples

Anonymized, formalin-fixed paraffin-embedded (FFPE) colon tissue from healthy individuals and IBD patients (including UC and CD patients) were acquired from ProteoGenex and Discovery Life Sciences under written informed consent. Patient information can be found in table 1.

### DSS colitis model

Mice were randomly assigned to 2 groups (n=15 mice/group). The first group was assigned as the control group, and the second group was assigned as the DSS-treated group. To induce colitis, mice were treated with 3.5% w/v Dextran sulfate sodium (DSS) in drinking water for 7 days (MP Biomedicals, 160110). DSS-treated mice were then switched to Innovive chlorinated water for 3 days and sacrificed at the 10-day time point. The control group had free access to drinking water throughout the ten-day course of the experiment. Mice were monitored for weight loss daily to assess disease progression. Weight loss and colon length were used as colitis readouts.

### Lipocalin-2 ELISA

Lipocalin-2 concentrations in fecal samples from control and DSS-treated mice were measured using a mouse lipocalin-2 Elisa kit (R&D Systems) according to the manufacturer’s protocol. Fecal pellets were homogenized in sterile PBS. 10 μl PBS were added per mg of stool. Homogenized fecal mixtures were centrifuged for 5 min at 2,000g at 4° C and supernatants were collected, and a 1:1000 dilution was measured by ELISA.

### Cytokine and Adipokine Array

For the cytokine and adipokine array, the mouse cytokine array panel A and the mouse adipokine array kit (R&D Systems, Minneapolis, MN, USA) were used. Mice were sacrificed (n=4 mice/group) by cervical dislocation, and the colons were removed from control and DSS-treated mice. The colons were placed in ice-cold PBS and cut open longitudinally. Luminal contents were removed by flushing with ice-cold PBS. The proximal, mid and distal colon were dissected as previously described^37^ and samples were snap frozen. Colon length varied from 60 to 80 mm. Tissue lysates were homogenized in 1% Triton X-100 in PBS containing a protease inhibitor cocktail (Roche Diagnostics, Mannheim, Germany) at 4° C. The lysates were frozen at -20° C, thawed, and centrifuged at 10,000 x *g* for 5 min to remove cellular debris. An aliquot of the sample was taken, and protein concentrations were quantified using the Pierce^TM^ BCA Protein Assay Kit (Thermofisher Scientific, USA). The tissue lysates (800 ug) were incubated with the cytokine and adipokine array membrane according to the manufacturer’s instructions with LI-COR detection. The arrays were scanned, and images were collected using the Li-Cor Odyssey^®^ CLx Imaging System (LI-COR Biosciences, USA). All measurements in this assay were conducted in duplicates. The mean signal intensity of each pair of duplicate spots was quantified using Image Studio Version 5.2. Mean signal intensities and statistical analysis performed are listed in table 2. To account for technical variability and to enable direct comparison across different blots, the integrated pixel densities were normalized using z-scores before plotting.

### Immunofluorescence staining and Image analysis

Mice were deeply anesthetized (2% sevoflurane) and transcardially perfused with ice-cold PBS followed by ice-cold 4% freshly depolymerized paraformaldehyde (PFA). Colon tissue was harvested and cryopreserved in sucrose and embedded in OCT. Sections (10 μm thick) were collected on plus charged slides and used for immunostaining. Frozen sections were thawed and washed 2 times with 10 mM Tris, pH 7.4, 0.9% NaCl, 0.05% Tween-20 (TBST) for 10 minutes at room temperature (RT), followed by one wash with PBS for 10 minutes at RT. Paraffin-embedded sections (5 μm thick) of human colon from healthy subjects and IBD patients were cleared in xylene and hydrated by passing through graded alcohols. Sections were subjected to heat-induced epitope retrieval (citrate buffer 0.1 M pH 6.0) using a pressure cooker at 110° C for 30 minutes. Sections were allowed to cool on ice for 45 minutes. After antigen retrieval, sections were washed in PBS for 5 minutes followed by two washes in TBST (10 minutes each). All sections were blocked in 10% donkey serum in TBST for 1 hour at RT to limit nonspecific antibody association. Incubation with primary antibodies was performed at 4° C in TBST plus 0.2% BSA for 48 to 72 hours. All antibodies used for immunofluorescence experiments are listed in Table 3. Sections were washed 4 times with PBS (10 minutes each time) and then incubated with secondary antibodies (see table 4) and 4’,6-diamidino-2-phenylindole (DAPI; Invitrogen, USA) to stain nuclei for 2 hours at RT. After 3 washes with PBS (20 minutes each time), sections were mounted using ProLong Diamond Antifade Mountant (Invitrogen, USA). Paraffin sections were treated with 10 mM CuSO4/50 mM NH4Cl solution for 5 minutes to quench tissue autofluorescence. Images were acquired using a Zeiss LSM 880 AiryScan inverted confocal microscope, with a Zeiss x20/0.8 NA or x40/1.4 NA Oil Plan-Apochromat DIC, (UV) VIS-IR (420762-9900) objective. Z-stacks with an interval of 0.5 μm were acquired and 3D images were rendered using 10.4.2 Imaris Software (Bitplane Inc.). At least 6 to 7 view fields at 40x magnification were imaged. For cell quantification of human colonic L-cells (Pyy+), we used the “surface” function and cells were automatically counted in Imaris. Count data was multiplied by 1 and then divided by the multiplied number of surveyed tissue areas (0.045 mm^2^) and view fields (6 to 7) to calculate the number of cells per 1 mm^2^, as previously described^38^. Every data point of a given graph corresponds to a single patient. For each patient we stained at least 2 FFPE colon sections. To measure the distance between L-cells (Pyy^+^), sensory neurons (Pgp9.5^+^), and T cells (CD3^+^), we used the “surface” function for each cell type, and we performed object to object statistics using Imaris. For relative frequency analysis of the distance between different cell types, we graphed all cells counted from each patient from Healthy individuals and IBD patients (see tables 5 and 6).

### Whole-mount intestine immunofluorescence

In brief, mice from control and DSS-treated groups were deeply anesthetized (2% sevoflurane) and transcardially perfused with ice-cold PBS followed by ice-cold 4% PFA. The colon was removed and placed in ice-cold PBS and cut open longitudinally. Luminal contents were flushed with ice-cold PBS and colon was placed in a 50 ml falcon tube containing 4% PFA overnight (O/N) at 4° C with gentle agitation. After washing the colon in PBS for 2 hours at 4° C, the proximal, mid and distal regions of the colon were dissected into specific sizes and locations as previously described^39^. Proximal colon size was between 20-24 mm, mid colon size between 30-33 mm and distal colon size between 18-23 mm. Proximal, mid and distal colon pieces (1 cm x 1 cm) were dehydrated in an ascending methanol series (50%, 80%,100%) for 5 minutes at 4° C. The permeabilization step included the incubation of colon pieces in 20% DMSO/methanol for 10 minutes RT, followed by rehydration in a descending methanol series (80%, 50%) and PBS for 10 minutes each RT with gentle agitation. Colon pieces were permeabilized with 0.2% Triton X-100 in PBS for 20 minutes RT, followed by incubation with Cytovista^TM^ antibody penetration buffer (Thermofisher, USA) for 1 hour at 4° C with gentle agitation. Colon pieces were then blocked with Cytovista^TM^ blocking buffer (Thermofisher, USA) O/N at 4° C with gentle agitation. Next, primary antibodies (see table 4) were diluted in Cytovista^TM^ antibody dilution buffer (Thermofisher, USA) and added to colon samples for three days at 4° C. After primary antibody incubation, the tissue was washed four times (30 min each) in Cytovista^TM^ wash buffer (Thermofisher, USA) and then incubated in Cytovista^TM^ antibody dilution buffer (Thermofisher, USA) with secondary antibodies (1:400) and DAPI (1:500) for 24 hours at 4° C (see table 5). Samples were again washed four times (30 min each) in Cytovista^TM^ wash buffer (Thermofisher, USA), dehydrated in an ascending methanol series (50%, 80%, 100%) at RT, then cleared with Cytovista^TM^ tissue clearing reagent (Thermofisher, USA) for 30 minutes at 4° C and mounted in Cytovista^TM^ tissue clearing enhancer (Thermofisher, USA) on slides. Slides were kept in the dark at 4° C until they were imaged.

### Microscopy and Image analysis

Whole-mount intestine samples were imaged on a Zeiss LSM 880 AiryScan inverted confocal microscope. At least 4 view fields (size 717226.1 μm^2^) at 20x magnification were imaged. Z-stacks with an interval of 2 μm were acquired and 3D images were rendered using 10.4.2 Imaris Software (Bitplane Inc.). Images were adjusted in gain and offset for optimal quality. Stack images were generated using the “volume rendering” function. 3D pictures were generated using the “snapshot” function. Final digital composites consisted of 2 μm z-slices, covering the full length of the colon mucosa, submucosal and myenteric plexus. For cell quantification of L-cells (Pyy^+^) and T cells (CD3^+^) we used the “surface” function and cells were automatically counted in Imaris. Cell counts were then divided by the size of the surveyed area (size 717226.1 μm^2^) and multiplied by 100,000 to calculate the number of cells per 0.1 mm^2^, as previously described^40^. Every data point of a given graph corresponds to a single animal. For measuring distance between L-cells (Pyy^+^), sensory neurons (Pgp9.5^+^) and T cells (CD3^+^), we used the “surface” function for each cell type, and we performed object to object statistics using Imaris. For relative frequency analysis of the distance between different cell types, we graphed all cells counted from each animal from control and DSS-treated mice (see tables 7 to 9).

### Human colonic epithelial cell atlas

An atlas of human colon epithelial cells was created by integrating healthy and IBD epithelial cells from colon samples from five previous scRNA-Seq studies^6,16,17,24,25^ using R packages Harmony (v1.2.0) and Seurat (v5.1.0). Cells and sample level annotations were curated to align metadata across the datasets, and epithelial cells were identified based on an epithelial gene signature (EPCAM, KRT8, KRT18) prior to integration. The cells in the integrated atlas were re-clustered into broad epithelial cell types and provided identities based on markers previously described^19,20^. Cluster 5, which comprised 90% of the EECs (as identified by the authors of the constituent datasets), was identified as EECs based on the high expression of markers CHGA and FEV, and further sub-clustered, identifying 4 subtypes – L type cells (PYY, GCG), Enterochromaffin cells (TPH1), Progenitors (NEUROG3), and D type cells (SST). Gene expression data were then aggregated at the sample level for each cell cluster, and pseudobulk differential expression analysis was conducted using the limma R package (v3.58.1). To identify overrepresented Gene Ontology (GO) terms within the genes differentially expressed L-cells from IBD samples, we performed an overrepresentation analysis using the PANTHER Classification System^41^.

### Quantification and Statistical Analysis

Data are presented as mean ± SEM. Comparisons between two conditions were analyzed by paired or unpaired 2-tailed student’s t-test or Mann-Whitney U test depending on normality of the data. The significance of the differences between more than two groups was evaluated using ANOVA followed by Turkey HSD post hoc testing or by Kruskal-Walli’s comparison test depending on the normality of the data. In figures, asterisks denote statistical significance (*p<0.05; **p<0.01; ***p<0.001; ****p<0.0001), unless indicated otherwise. Experiments were repeated at least twice. Differences were considered to be statistically significant when p<0.05. Statistical analysis was performed in GraphPad Prism 10.4.1.

## Author Contributions

These investigations were conducted, supporting methodologies developed, and data were acquired and validated by IAG. The Human colonic epithelial cell atlas and meta-analysis of published scRNA-Seq data was performed by PKV. Data were visualized by IAG and PKV, and the original draft was prepared by IAG. The project was conceptualized, data were reviewed, and the manuscript was reviewed and edited by IAG, PKV, TW, and RG. This work was funded by Pfizer Inc. and supported by Pfizer’s WRDM Postdoctoral Fellowship Program.

## Acknowledgments

We would like to acknowledge all the members of the Immune Homeostasis Group for all their expert input and feedback on this project. We would like also to acknowledge and thank Dr. Kendra Bence, Dr. Ellene Mashalidis, and Dr. Michelle Rooks for their feedback and valuable input on this project as members of IAG’s Postdoctoral Fellow committee. We would also like to thank Pfizer’s Optical Microscopy Center and more specifically Dr. Shoh Asano for his technical support and microscopy expertise. We gratefully acknowledge Dr. Jie Quan and Dr. Fridrik J. Karlsson for their expert input in the construction and interpretation of the human colonic epithelial cell atlas.

