## Supplemental Materials for "L-Cells are the Functional Neuropod Cell in Human Gastrointestinal Tract and are Dysregulated in Inflammatory Bowel Disease (IBD)"

The PDF file includes:

Supplementary Figures 1 to 9

Tables 1 to 9

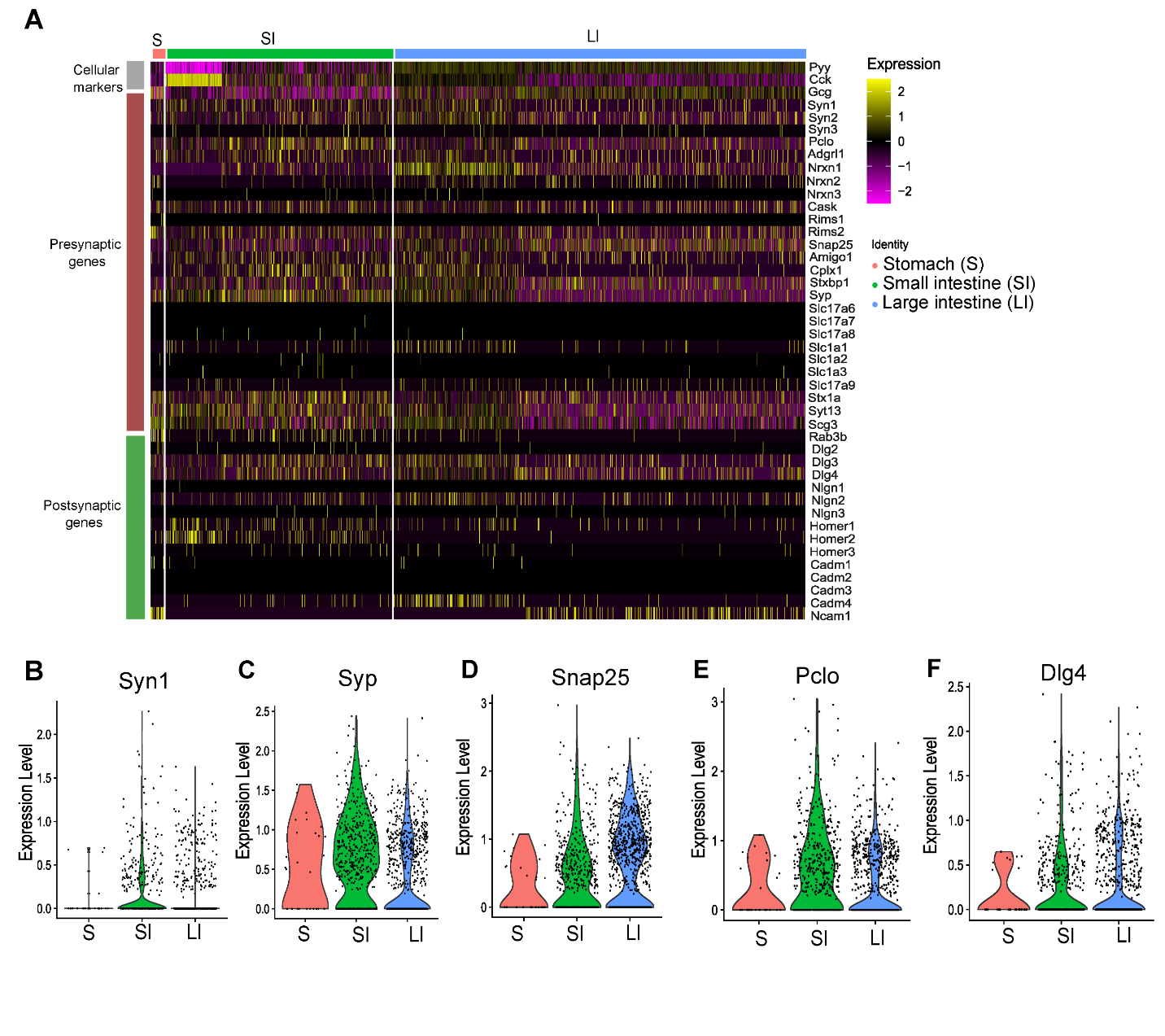

**Supplementary Figure 1**. **L-cells along the murine colon express pre- and postsynaptic genes.**

(A) Normalized expression of a select list of pre- and postsynaptic genes in L-cells from mouse stomach (S), small intestine (SI), and large intestine (LI).

(B-F) Gene expression of *Syn1, Syp, Snal25, Pclo,* and *Dlg4* in L-cells.

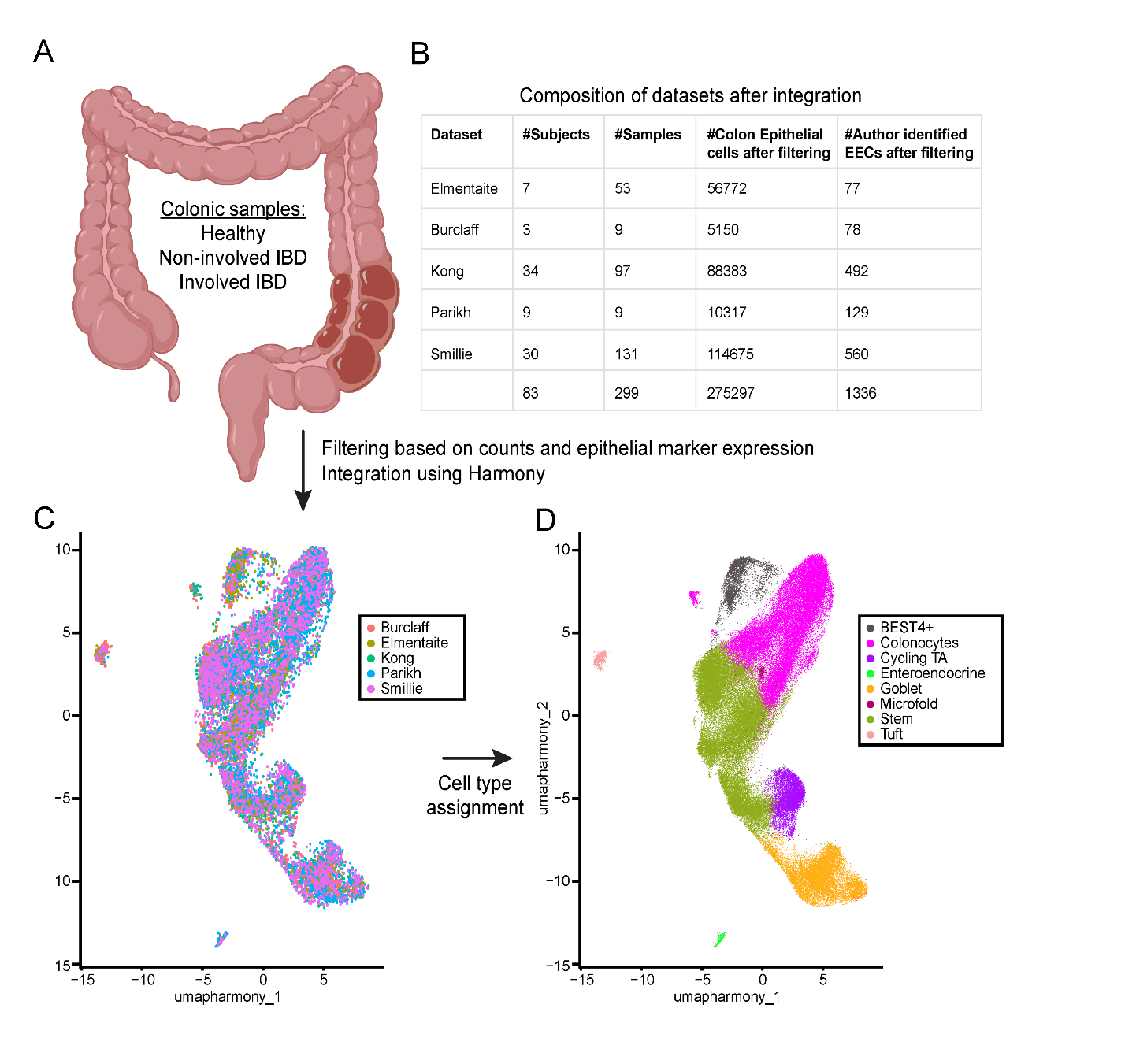

**Supplementary Figure 2. Computational workflow analysis of scRNA-Seq IBD samples from several datasets.**	(A) Schematic illustration depicting the human colon. Biopsies were taken from Healthy individuals, non-involved (sites with no significant inflammation from active IBD patients) and involved IBD (sites with significant inflammation from active IBD patients).

(B) Table depicting the datasets and the numbers of subjects, samples, colon epithelial cells after filtering and the author identified EECs after filtering.

(C) UMAP of atlas showing cells from the five datasets after Harmony integration.

(D) UMAP of atlas post cell type assignment.

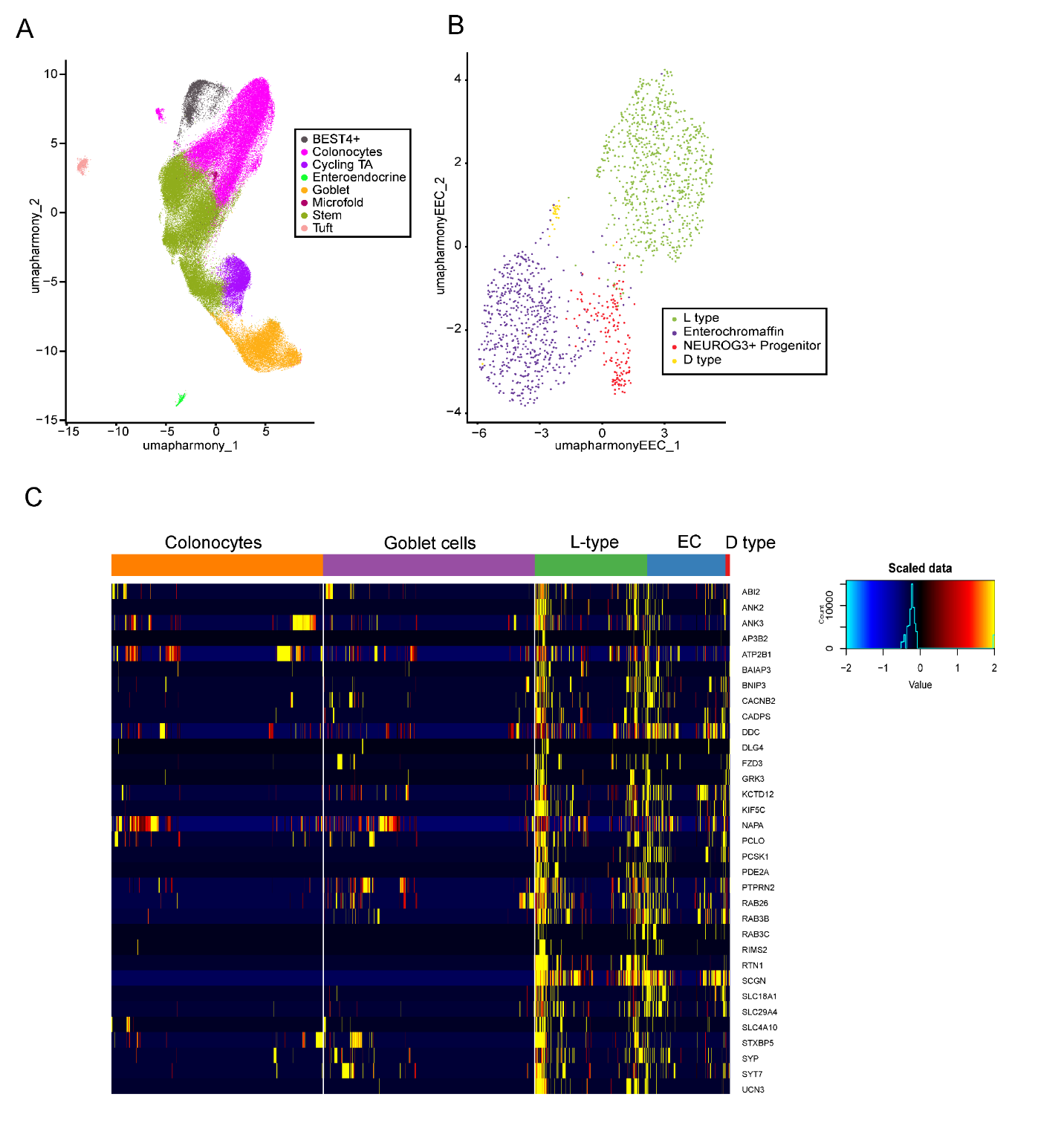
**Supplementary Figure 3.** **L-cells in the healthy human colon express pre- and postsynaptic genes.**

(A) UMAP of the human colon epithelial cell atlas compiled in this work.

(B) UMAP of the subsets of enteroendocrine cells identified in the compiled epithelial cell atlas.

(C) Normalized expression of select pre- and postsynaptic genes in representative cell populations of colonocytes, goblet cells, L-cells, enterochromaffin cells (EC), and D-type cells in the human colon.

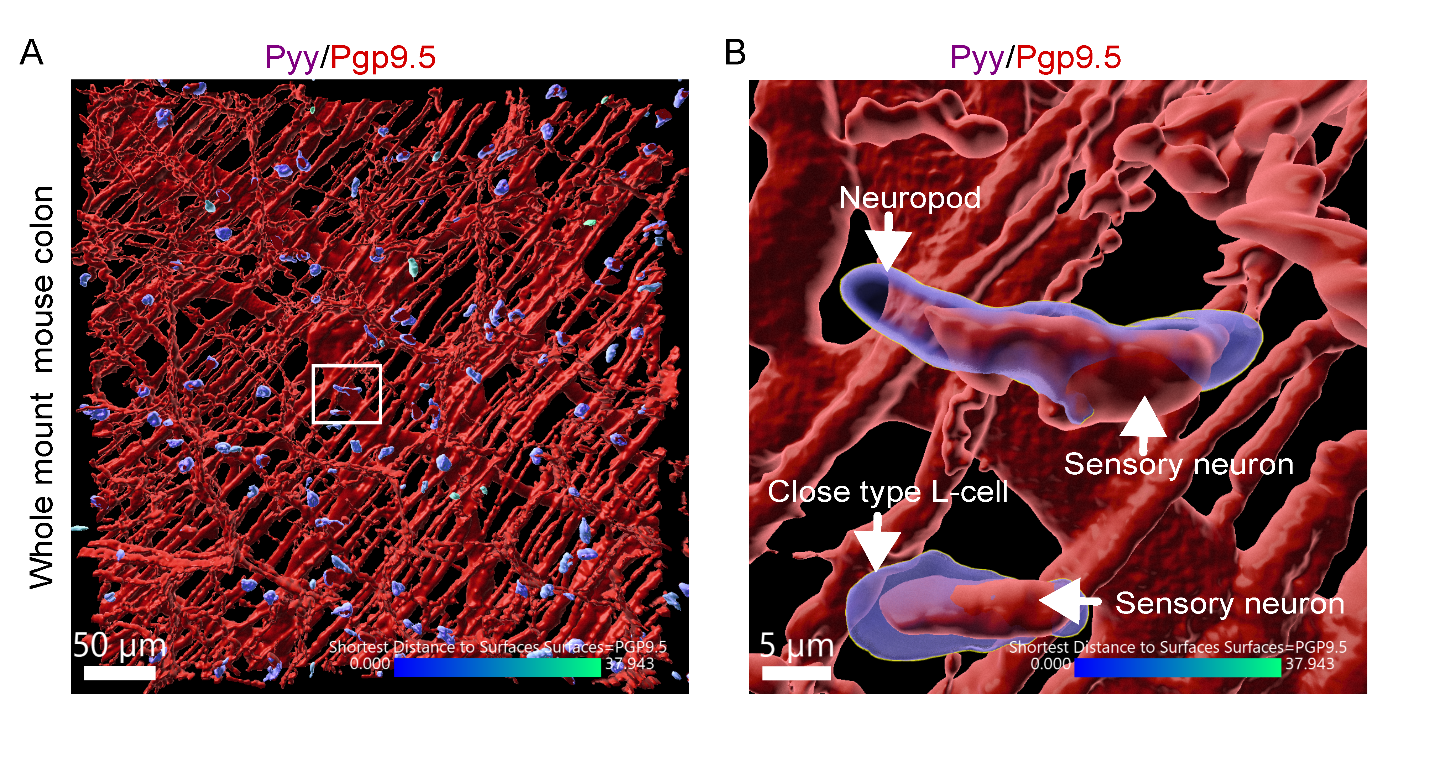
**Supplementary Figure 4. Neuropod cells and close type L-cells are proximal to sensory neurons.**

(A) Representative confocal image of whole mount mouse colon stained for Pyy and Pgp9.5. Scale bar 50 μm.

(B) Insert: Magnified confocal image depicting proximity of Pyy^+^ cells with neuropod and non-neuropod morphology (close type L-cell) to Pgp9.5^+^ sensory neurons. Scale bar 5 μm.

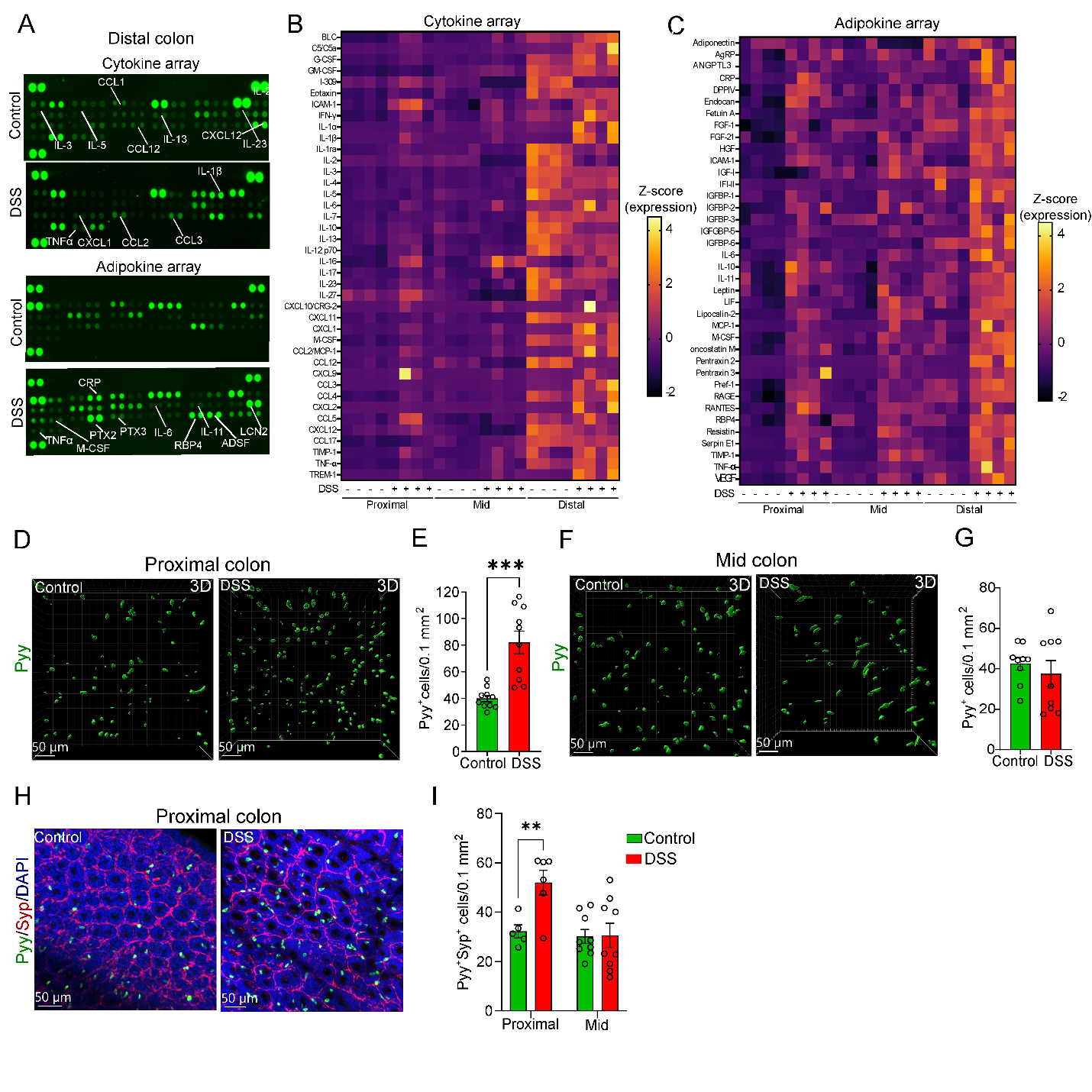

**Supplementary Figure 5.** **Acute DSS colitis induces spatially distinct effects on the abundance of neuropod cells along the colon epithelium.**

(A) Mouse cytokine and adipokine array analysis of distal colon tissue lysates (800 ug) from control and DSS-treated mice.

(B-C) Quantitation of tissue cytokines, chemokines and adipokines in control and DSS-treated mice. Heatmap illustrations of the 38 mouse adipokines array and 40 mouse cytokines/chemokines array (n=4 mice per group analyzed in duplicate). Signals from control mice normalized to 1 and compared with those from DSS-treated mice (upregulated cytokines/chemokines and adipokine proteins in yellow, downregulated in magenta).

(D) Representative 3D images of whole mount proximal colon stained for Pyy of control and DSS-treated mice. Scale bar 50 μm.

(E) Quantification of Pyy^+^ cells per 0.1 mm^2^ in the proximal colon of control (n=10) and DSS-treated (n=10) mice.

(F) Representative 3D images of whole mount mid colon stained for Pyy of control and DSS-treated mice. Scale bar 50 μm.

(G) Quantification of Pyy^+^ cells per 0.1 mm^2^ in the mid colon of control (n=9) and DSS-treated (n=8) mice.

(H) Representative confocal images of whole mount proximal colon stained for Pyy and Syp in control and DSS-treated mice. DAPI, Scale bars, 50 μm.	(I) Quantification of Pyy^+^Syp^+^ cells per 0.1 mm^2^ in the proximal and mid colon of control and DSS-treated mice (n=5-9 mice/group).

Plotted are means ± SEM. *p<0.05, **p<0.01, ***p<0.001, ****p<0.000 by unpaired two-tailed tests. Experiments were repeated at least twice.

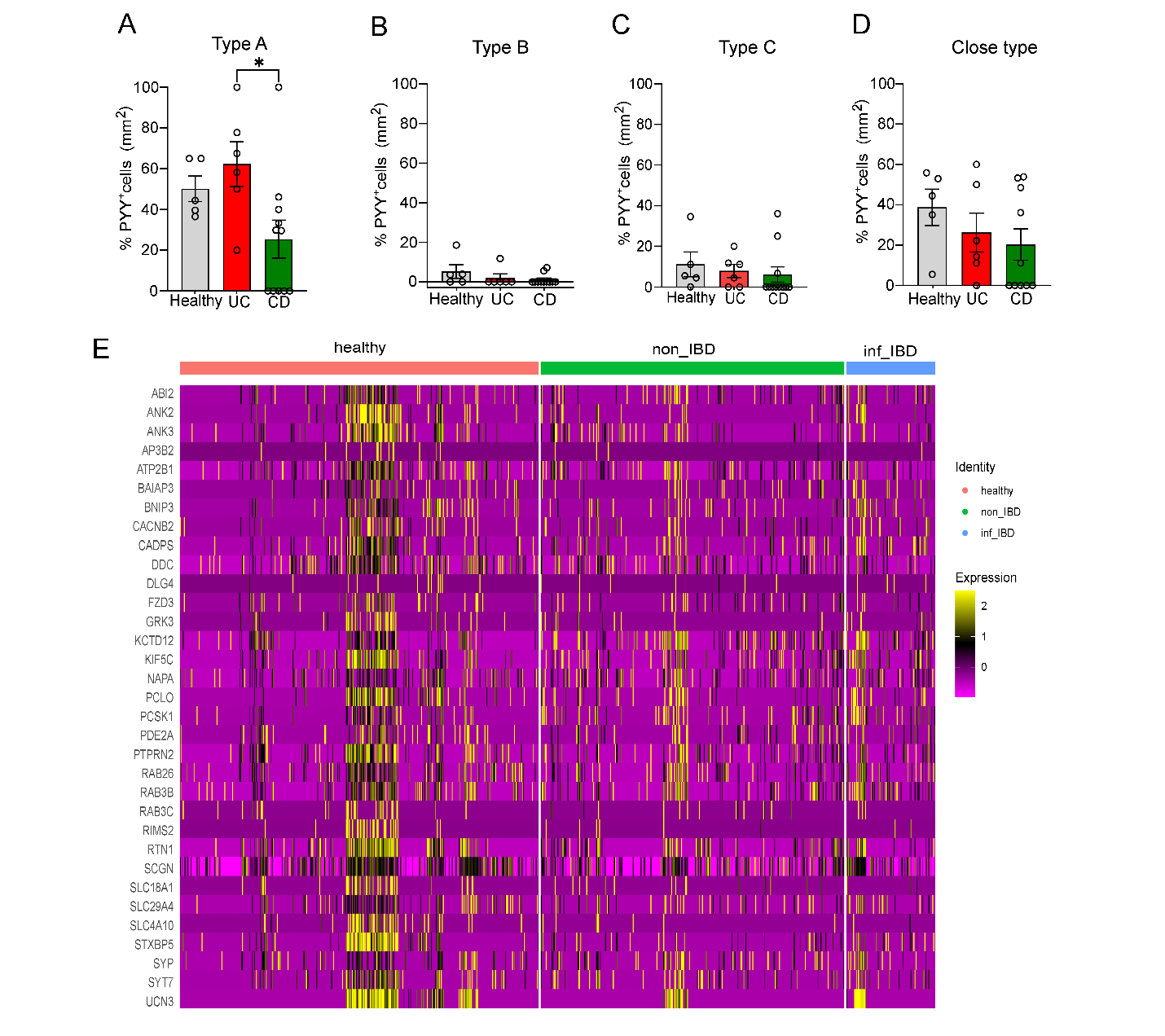

**Supplementary Figure 6. Expression of pre-and postsynaptic genes in colonic L-cells of healthy subjects and IBD patients.**

(A-D) Percentage of the different types of Pyy^+^ cells, normalized to total number of Pyy^+^ cells per mm^2^ in colon of healthy subjects (n=5), UC (n=6) and CD (n=11) patients.

(E) Normalized expression of select pre- and postsynaptic genes in L-cells from the colon of healthy subjects, and non-involved (non_IBD) and involved (inf_IBD) samples from active IBD patients.

Plotted are means ± SEM. *p<0.05 by ANOVA followed by Kruskal-Wallis comparison test.

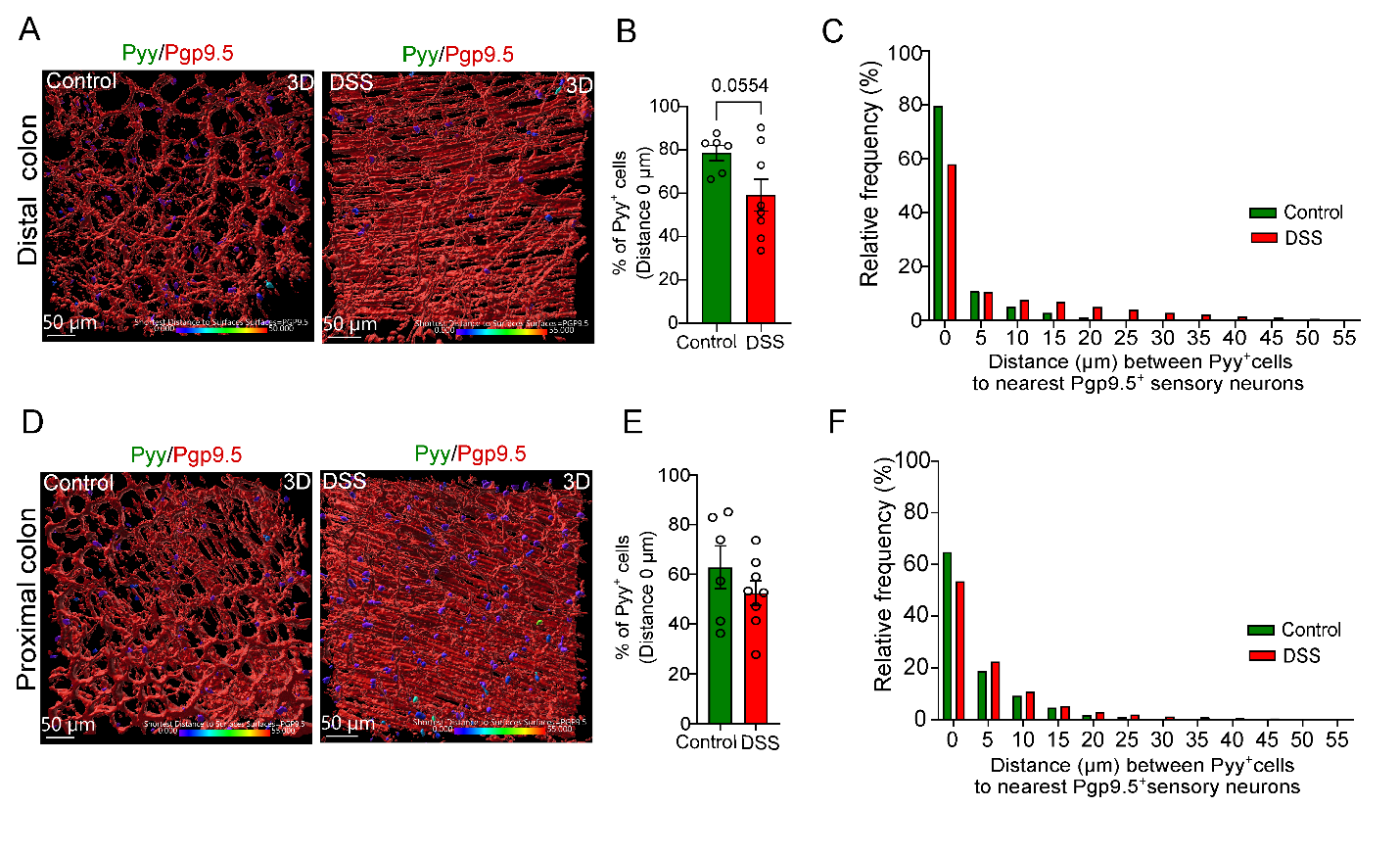

**Supplementary Figure 7*.* Acute DSS colitis did not alter significantly the proximity of sensory neurons to neuropod cells in the distal and proximal colon.**

(A) Representative 3D image of whole mount distal colon from control and DSS-treated mice stained for Pyy and Pgp9.5. Scale bar 50 μm.

(B) Percentage of Pyy^+^ cells in proximity (Distance 0 μm) to Pgp9.5+ sensory neurons in the distal colon of control (n= 6) and DSS-treated (n=8) mice.

(C) Relative frequency of distance (μm) between Pyy^+^ cells and nearest Pgp9.5^+^ssensory neurons in the distal colon of control (pooled data from 6 mice) and DSS-treated mice (pooled data from 8 mice).

(D) Representative 3D image of whole mount proximal colon from control and DSS-treated mice stained for Pyy and Pgp9.5. Scale bar 50 μm.

(E) Percentage of Pyy^+^ cells in proximity (Distance 0 μm) to Pgp9.5+ sensory neurons in the proximal colon of control (n= 6) and DSS-treated (n=7) mice.

(F) Relative frequency of distance (μm) between Pyy^+^ cells and nearest Pgp9.5^+^sensory neurons in the proximal colon of control (pooled data from 6 mice) and DSS-treated mice (pooled data from 7 mice).

Plotted are means ± SEM. p>0.05 by unpaired two-tailed tests. Experiments were repeated at least twice.

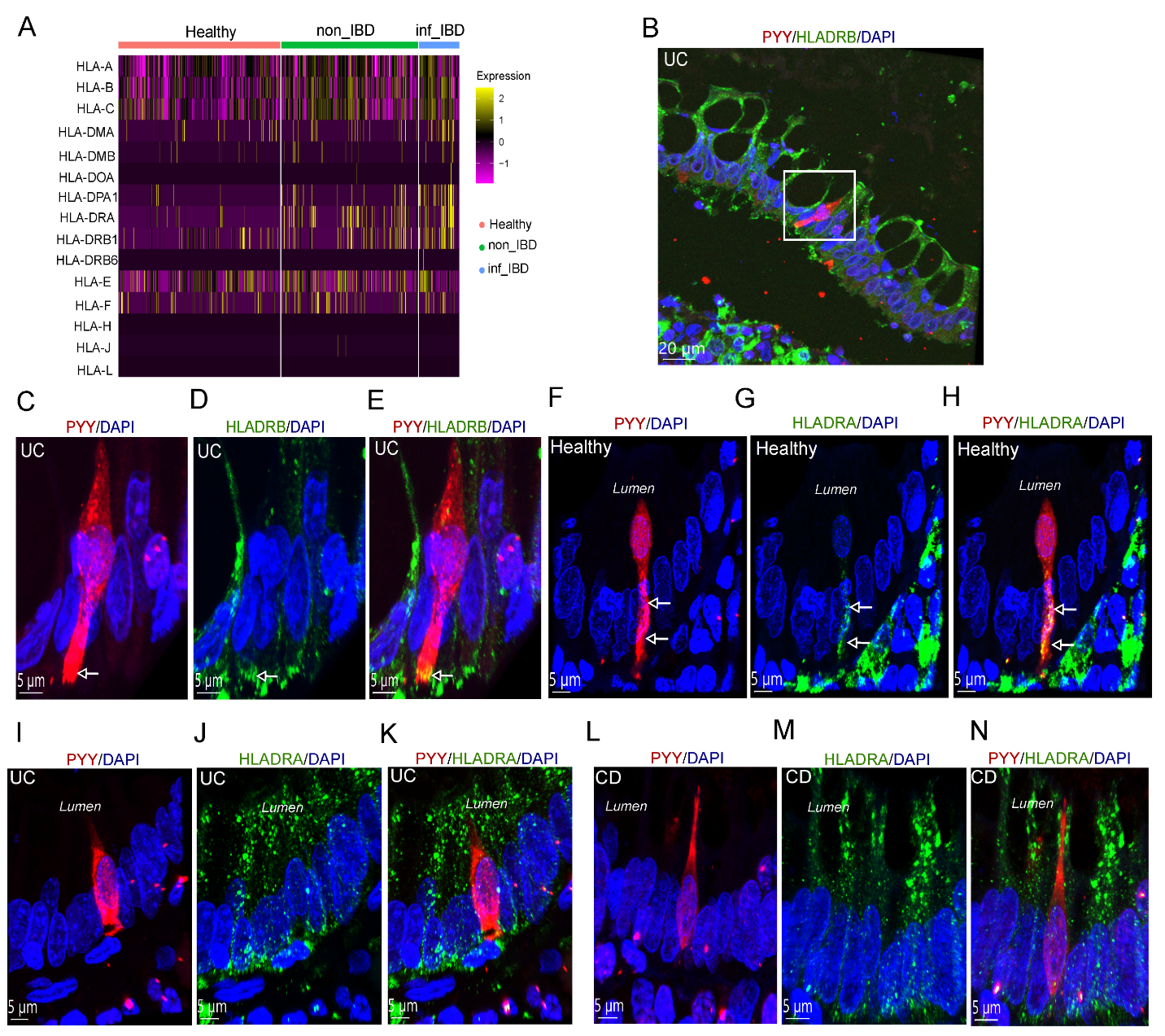

**Supplementary Figure 8. Increased expression of genes associated with antigen presentation in colonic L-cells of IBD patients.**

(A) Normalized expression of a select list of HLA family genes in L-cells in the colon of healthy subjects, and non-involved (non_IBD) and involved (inf_IBD) samples from active IBD patients.

(B-E) Representative confocal image of a 5 µm thick FFPE human colon section of a UC patient stained for PYY and HLA-DRB. Scale bar 20 μm. Inset: Representative magnified confocal images from the colon of a UC patient stained for PYY and HLA-DRB; arrows depict colocalization of PYY and HLA-DRB. Scale bar 5 μm.

(F-H) Representative magnified confocal images from the colon of a healthy subject stained for PYY and HLA-DRA; arrows depict colocalization of PYY and HLA-DRA. Scale bar 5 μm.

(I-K) Representative magnified confocal images from the colon of a UC patient stained for PYY and HLA-DRA; no colocalization is evident for PYY and HLA-DRA. Scale bar 5 μm.

(L-N) Representative magnified confocal images from the colon of a CD patient stained for PYY and HLA-DRA; no colocalization is evident for PYY and HLA-DRA. Scale bar 5 μm.

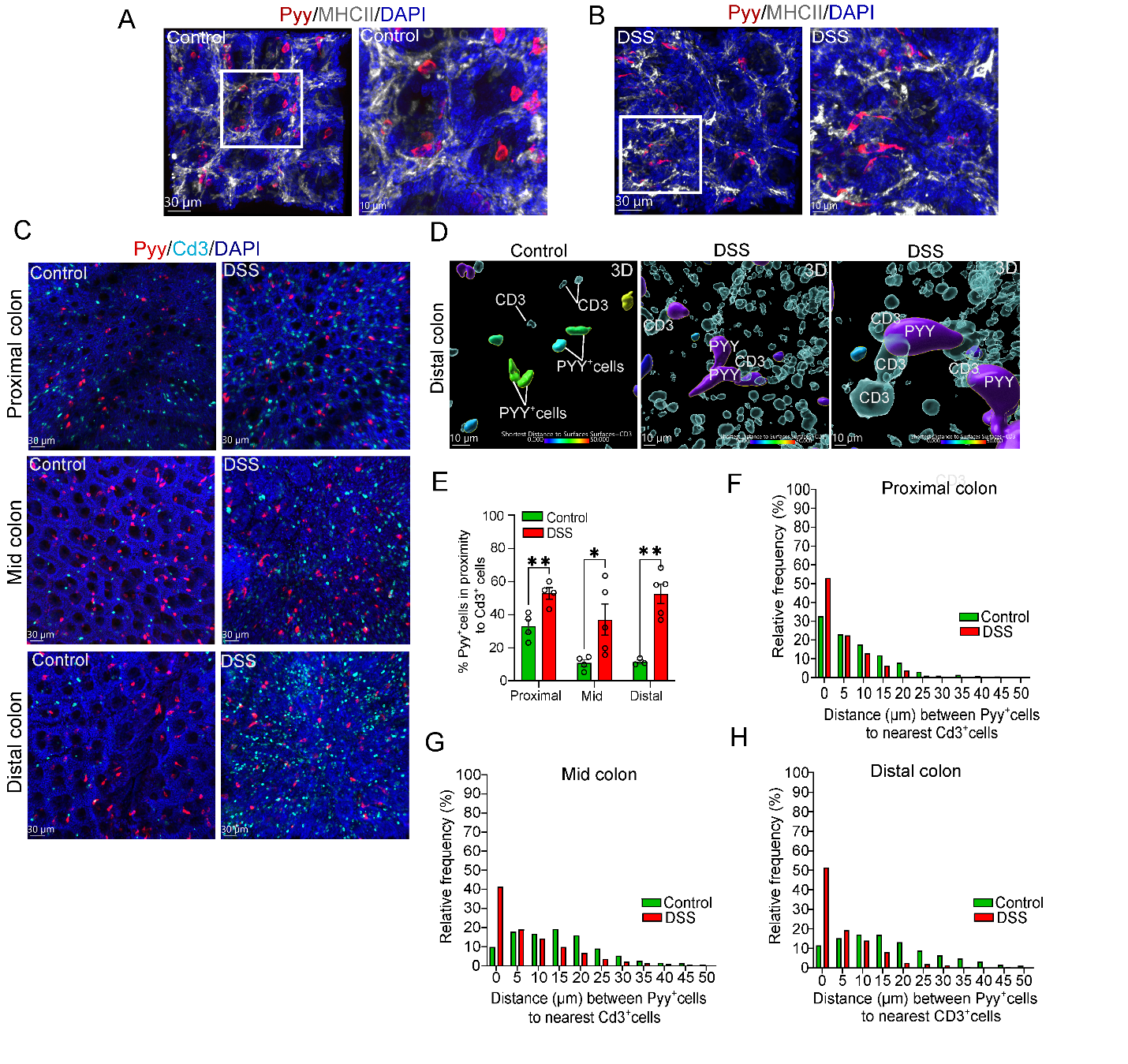
**Supplementary Figure 9. Acute DSS colitis increases the L-cell to T cells proximity in the different regions of the colon.**

(A-B) Representative confocal images of whole mount mouse colon from control and DSS-treated mice stained for Pyy and MHCII. 3D images depict the proximity of Pyy^+^ cells to MHCII^+^ cells. MHCII is expressed in the epithelium and lamina propria in mice with DSS colitis. Scale bar 30 μm.

(C) Representative confocal images of whole mount mouse proximal, mid and distal colon from control and DSS-treated mice stained for Pyy and CD3. Scale bar 30 μm.

(D) Magnified representative 3D images of whole mount distal colon stained for Pyy and CD3 of control and DSS-treated mice. Scale bar 10 μm

(E) Percentage of Pyy^+^ cells in proximity (Distance 0 μm) to CD3^+^ T cells in the proximal, mid and distal colon of control and DSS-treated mice (n=3-5 mice/group).

(F) Relative frequency of distance (μm) between Pyy^+^ cells and nearest CD3^+^ T cells in the proximal colon of control (pooled data from 4 mice) and DSS-treated mice (pooled data from 4 mice).

(G) Relative frequency of distance (μm) between Pyy^+^ cells and nearest CD3^+^ T cells in the mid colon of control (pooled data from 4 mice) and DSS-treated mice (pooled data from 5 mice).

(H) Relative frequency of distance (μm) between Pyy^+^ cells and nearest CD3^+^ T cells in the distal colon of control (pooled data from 3 mice) and DSS-treated mice (pooled data from 5 mice).

Plotted are means ± SEM. *p<0.05, **p<0.01 by unpaired two-tailed tests.

**Table 1.** Patient characteristics-Tissue samples

| **Patient ID#** | **Vendor** | **Age** | **Race** | **Sex** | **Tissue** | **Diagnosis** | **Treatment** | **Tumor %** | **Necrosis %** |
| --- | --- | --- | --- | --- | --- | --- | --- | --- | --- |
| 1 | ProteoGenex | 60 | Caucasian | F | Colon | Normal tissue | N/A | 0 | 0 |
| 2 | ProteoGenex | 53 | Caucasian | F | Colon | Normal tissue | N/A | 0 | 0 |
| 3 | ProteoGenex | 62 | Caucasian | F | Colon | Normal tissue | N/A | 0 | 0 |
| 4 | ProteoGenex | 28 | Caucasian | M | Colon | Normal tissue | N/A | 0 | 0 |
| 5 | ProteoGenex | 71 | Caucasian | F | Colon | Normal tissue | N/A | 0 | 0 |
| 6 | Discovery Life Sciences | 65 | White | M | Colon | Ulcerative colitis | Active Tx | 0 | 0 |
| 7 | Discovery Life Sciences | 51 | White | F | Colon | Ulcerative colitis | Active Tx | 0 | 0 |
| 8 | Discovery Life Sciences | 21 | White | M | Colon | Ulcerative colitis | Pre Tx | 0 | 0 |
| 9 | Discovery Life Sciences | 40 | White | M | Colon | Ulcerative colitis | Post Tx | 0 | 0 |
| 10 | Discovery Life Sciences | 22 | White | F | Colon | Ulcerative colitis | Pre Tx | 0 | 0 |
| 11 | Discovery Life Sciences | 32 | White | F | Colon | Ulcerative colitis | Pre Tx | 0 | 0 |
| 12 | Discovery Life Sciences | N/A | White | F | Colon | Crohn’s disease | Active Tx | 0 | 0 |
| 13 | Discovery Life Sciences | 37 | White | M | Colon | Crohn’s Disease | Pre Tx | 0 | 0 |
| 14 | Discovery Life Sciences | 70 | White | F | Colon | Crohn’s Disease | Pre Tx | 0 | 0 |
| 15 | Discovery Life Sciences | 58 | White | M | Colon | Crohn’s Disease | Post Tx | 0 | 0 |
| 16 | Discovery Life Sciences | 55 | White | F | Colon | Crohn’s Disease | Post Tx | 0 | 0 |
| 17 | Discovery Life Sciences | 41 | White | M | Colon | Crohn’s Disease | Post Tx | 0 | 0 |
| 18 | Discovery Life Sciences | 19 | White | M | Colon, Ascending | Crohn’s Disease | Pre Tx | 0 | 0 |
| 19 | Discovery Life Sciences | 20 | White | M | Colon, Ascending | Crohn’s Disease | Pre Tx | 0 | 0 |
| 20 | Discovery Life Sciences | 36 | White | F | Colon, Rectosigmoid | Crohn’s Disease | Pre Tx | 0 | 0 |
| 21 | Discovery Life Sciences | 41 | White | F | Colon | Crohn’s Disease | Post Tx | 0 | 0 |
| 22 | Discovery Life Sciences | 32 | White | M | Colon | Crohn’s Disease | Pre Tx | 0 | 0 |

Tx: Treatment with x medication not specified

**Table 2.** Cytokine and Adipokine array for distal colon.

| **Analytes** | **Mean Intensity Control**  **(n=4)** | **Mean Intensity**  **DSS (Day 10) (n=4)** | **Difference** | **SE of difference** | **P-value** | **Multiple T-test** |
| --- | --- | --- | --- | --- | --- | --- |
| **IL-1β** | 13321 | 141446 | -128124 | 39895 | 0.0183 | Unpaired t-test |
| **IFN-γ** | 19816 | 30971 | -11154 | 5652 | 0.0959 | Unpaired t-test |
| **IL-1a** | 26029 | 86871 | -60842 | 28190 | 0.0743 | Unpaired t-test |
| **IL-2** | 20016 | 7238 | 12778 | 2027 | 0.0007 | Unpaired t-test |
| **IL-3** | 12441 | 9458 | 2983 | 814.3 | 0.0105 | Unpaired t-test |
| **IL-4** | 16929 | 14621 | 2308 | 1885 | 0.2668 | Unpaired t-test |
| **IL-5** | 5501 | 3215 | 2287 | 806.7 | 0.0298 | Unpaired t-test |
| **IL-6** | 5534 | 11181 | -5647 | 2256 | 0.0463 | Unpaired t-test |
| **IL-7** | 17316 | 15346 | 1971 | 1626 | 0.2711 | Unpaired t-test |
| **IL-10** | 16354 | 16011 | 342.3 | 2010 | 0.8704 | Unpaired t-test |
| **IL-11** | 15501 | 24491 | -8990 | 2434 | 0.0102 | Unpaired t-test |
| **OSM** | 12688 | 22403 | -9715 | 1909 | 0.0022 | Unpaired t-test |
| **IL-13** | 16529 | 11721 | 4808 | 1698 | 0.0299 | Unpaired t-test |
| **IL-12p70** | 6580 | 5295 | 1286 | 892.8 | 0.20 | Unpaired t-test |
| **IL-16** | 51404 | 84271 | -32867 | 15674 | 0.0808 | Unpaired t-test |
| **IL-17** | 21716 | 19883 | 1833 | 4181 | 0.6764 | Unpaired t-test |
| **IL-23** | 26904 | 15633 | 11271 | 3975 | 0.0297 | Unpaired t-test |
| **IL-27** | 43304 | 25321 | 17983 | 9150 | 0.097 | Unpaired t-test |
| **M-CSF** | 18701 | 35216 | -16515 | 3697 | 0.0043 | Unpaired t-test |
| **GM-CSF** | 15316 | 14883 | 433 | 1520 | 0.7853 | Unpaired t-test |
| **G-CSF** | 15604 | 20396 | -4792 | 3257 | 0.1916 | Unpaired t-test |
| **TNF-α** | 13994 | 20942 | -6948 | 2410 | 0.028 | Unpaired t-test |
| **CCL1** | 29541 | 10933 | 18608 | 2010 | <0.0001 | Unpaired t-test |
| **CXCL1** | 6936 | 6372 | 564.3 | 683 | 0.4403 | Unpaired t-test |
| **CXCL10** | 18516 | 78271 | -59754 | 47211 | 0.0286 | Mann-Whitney |
| **CXCL11** | 15791 | 12321 | 3471 | 1663 | 0.0819 | Unpaired t-test |
| **CCL17** | 7474 | 6493 | 980.5 | 383.1 | 0.0429 | Unpaired t-test |
| **CXCL1** | 6915 | 21837 | -14922 | 5248 | 0.0294 | Unpaired t-test |
| **CCL2** | 17091 | 63883 | -46792 | 16706 | 0.0311 | Unpaired t-test |
| **CCL12** | 22879 | 9876 | 13003 | 1557 | 0.0002 | Unpaired t-test |
| **CXCL9** | 15616 | 31233 | -15617 | 11559 | 0.2254 | Unpaired t-test |
| **CXCL13** | 24916 | 34533 | -9617 | 4102 | 0.0575 | Unpaired t-test |
| **CCL3** | 12150 | 48958 | -36808 | 12075 | 0.0226 | Unpaired t-test |
| **CCL4** | 10799 | 12141 | -1342 | 2487 | 0.6089 | Unpaired t-test |
| **CXCL12** | 178716 | 115433 | 63283 | 15586 | 0.0066 | Unpaired t-test |
| **CCL5** | 16529 | 17608 | -1079 | 3655 | 0.7777 | Unpaired t-test |
| **TREM-1** | 13179 | 85621 | -72442 | 20402 | 0.0121 | Unpaired t-test |
| **TIMP1** | 81072 | 181131 | -100058 | 53244 | 0.1093 | Unpaired t-test |
| **IL-1ra** | 2430966 | 909096 | 1521870 | 337826 | 0.0041 | Unpaired t-test |
| **C5/C5a** | 90441 | 379346 | -288904 | 109939 | 0.0392 | Unpaired t-test |
| **ICAM1** | 795716 | 1329596 | -533879 | 160737 | 0.016 | Unpaired t-test |
| **LCN2** | 52988 | 1598391 | -1545402 | 49544 | <0.0001 | Unpaired t-test |
| **RBP4** | 230351 | 353766 | -123415 | 25169 | 0.0027 | Unpaired t-test |
| **OB** | 9595 | 13641 | -4046 | 1326 | 0.0225 | Unpaired t-test |
| **ADSF** | 55938 | 351516 | -295577 | 30305 | <0.0001 | Unpaired t-test |
| **CRP** | 26913 | 165953 | -139040 | 38265 | 0.0109 | Unpaired t-test |
| **PTX2** | 36538 | 1539016 | -1502477 | 370277 | 0.0067 | Unpaired t-test |
| **PTX3** | 8478 | 41141 | -32662 | 5821 | 0.0014 | Unpaired t-test |
| **AgRP** | 33601 | 37366 | -3765 | 11299 | 0.7503 | Unpaired t-test |
| **ANGPTL3** | 10980 | 25453 | -14473 | 4244 | 0.0143 | Unpaired t-test |
| **DPPIV** | 120201 | 151766 | -31565 | 21002 | 0.1836 | Unpaired t-test |
| **ESM-1** | 12145 | 15366 | -3221 | 1603 | 0.0912 | Unpaired t-test |
| **FGF-1** | 241476 | 246266 | -4790 | 19214 | 0.8115 | Unpaired t-test |
| **FGF-21** | 8340 | 11553 | -3213 | 833.8 | 0.0084 | Unpaired t-test |
| **HGF** | 15426 | 52303 | -36877 | 6977 | 0.0019 | Unpaired t-test |
| **Acrp30** | 2242226 | 2157141 | 85085 | 420976 | 0.8465 | Unpaired t-test |
| **AHSG** | 375726 | 543266 | -167540 | 28324 | 0.001 | Unpaired t-test |
| **PAI-1** | 14106 | 73191 | -59085 | 15350 | 0.031 | Unpaired t-test |
| **RAGE** | 8142 | 10528 | -2386 | 708.6 | 0.0151 | Unpaired t-test |
| **LIF** | 10257 | 18828 | -8571 | 2695 | 0.0191 | Unpaired t-test |
| **VEGF** | 12487 | 17628 | -5141 | 3017 | 0.1393 | Unpaired t-test |
| **Pref-1** | 9211 | 13566 | -4355 | 1068 | 0.0065 | Unpaired t-test |
| **IGF-I** | 20926 | 19178 | 1748 | 2065 | 0.4297 | Unpaired t-test |
| **IGF-II** | 32388 | 65378 | -32990 | 18981 | 0.1329 | Unpaired t-test |
| **IGFBP-1** | 9972 | 23466 | -13493 | 3588 | 0.0094 | Unpaired t-test |
| **IGFBP-2** | 81788 | 155766 | -73977 | 27987 | 0.0384 | Unpaired t-test |
| **IGFBP-3** | 65938 | 108678 | -42740 | 23305 | 0.1164 | Unpaired t-test |
| **IGFBP-5** | 19701 | 102766 | -83065 | 16498 | 0.0024 | Unpaired t-test |
| **IGFBP-6** | 94901 | 106803 | -11902 | 16405 | 0.4955 | Unpaired t-test |

**Table 3.** Primary antibodies

| **Target** | **Source** | **Catalog#** | **Company** | **Dilution** | **Target Species** |
| --- | --- | --- | --- | --- | --- |
| Pyy | Guinea pig | 16066 | Progen | 1:500 in frozen and paraffin sections; 1:2000 in whole mount colon | Mouse, Human |
| Syn1 | Rabbit | 5297S | Cell Signaling | 1:100 in frozen and paraffin sections | Mouse |
| Syp | Rabbit | 17785-1-AP | Proteintech | 1:100 in frozen and paraffin sections; 1:1000 in whole mount colon | Mouse, Human |
| Syp-CoraLite® Plus 647 | Rabbit | CL647-17785 | Proteintech | 1:100 in frozen and paraffin sections | Mouse, Human |
| UCH-L1/Pgp9.5 | Rabbit | 14730-1-AP | Proteintech | 1:100 in frozen and paraffin sections; 1:1000 in whole mount colon | Mouse,  Human |
| UCH-L1/Pgp9.5- CoraLite® 594 | Rabbit | CL594-14730 | Proteintech | 1:100 in frozen and paraffin sections | Mouse |
| CD3 | Rat | Ab11089 | Abcam | 1:100 in frozen and paraffin sections; 1:500 in whole mount colon | Mouse, Human |
| HLA-DRB1 | Rabbit | 15862-1-AP | Proteintech | 1:100 in paraffin sections | Human |
| HLA-DRA | Rabbit | 97971S | Cell Signaling | 1:100 in paraffin sections | Human |
| MHCII | Rat | 55699 | BD Bioscience | 1:1000 in whole mount colon | Mouse |

**Table 4.** Secondary antibodies

| **Target** | **Source** | **Fluor** | **Catalog#** | **Company** | **Dilution** |
| --- | --- | --- | --- | --- | --- |
| Rabbit IgG | Donkey | AF568 | A10042 | Thermofisher | 1:400-1:500 |
| Rabbit IgG | Donkey | AF488 | A-21206 | Thermofisher | 1:400-1:500 |
| Rabbit IgG | Donkey | AF647 | A-31573 | Thermofisher | 1:400-1:500 |
| Rat IgG | Donkey | AF647 | A78947 | Thermofisher | 1:400-1:500 |
| Goat IgG | Donkey | AF647 | A-21447 | Thermofisher | 1:400-1:500 |
| Guinea pig IgG | Donkey | AF568 | 706-575-148 | Jackson Laboratories | 1:800-1:1000 |

**Table 5.** Frequency distribution of the distance between Pyy^+^ cells and Pgp9.5^+^ sensory neurons in the colon of Healthy individuals and IBD patients.

| Distance (μm) | Healthy | UC | CD |
| --- | --- | --- | --- |
| 0 | 48.03 | 15.85 | 16.92 |
| 5 | 26.47 | 23.17 | 33.84 |
| 10 | 16.66 | 15.85 | 26.15 |
| 15 | 5.88 | 12.19 | 12.30 |
| 20 | 1.96 | 9.75 | 4.61 |
| 25 | 0.98 | 9.756 | 1.53 |
| 30 | 0 | 3.65 | 1.53 |
| 35 | 0 | 3.65 | 1.53 |
| 40 | 0 | 1.219 | 0 |
| 45 | 0 | 1.219 | 1.53 |
| 50 | 0 | 2.43 | 0 |
| 55 | 0 | 1.219 | 0 |
| Number of pooled cells | **102** | **83** | **65** |

**Table 6.** Frequency distribution of the distance between Pyy^+^ cells and CD3^+^ T cells in the colon of Healthy individuals and IBD patients

| Distance (μm) | Healthy | UC | CD |
| --- | --- | --- | --- |
| 0 | 35.21 | 40.27 | 44.59 |
| 5 | 25.35 | 15.27 | 22.97 |
| 10 | 9.85 | 20.83 | 16.21 |
| 15 | 5.63 | 11.11 | 10.81 |
| 20 | 11.26 | 5.55 | 0 |
| 25 | 7.04 | 2.77 | 2.70 |
| 30 | 5.63 | 4.16 | 2.70 |
| Number of pooled cells | **78** | **72** | **81** |

**Table 7**. Frequency distribution of the distance between Pyy^+^ cells and Pgp9.5^+^ sensory neurons in the various parts of the healthy mouse colon.

| Distance (μm) | Proximal colon | Mid colon | Distal colon |
| --- | --- | --- | --- |
| 0 | 72.00 | 78.98 | 81.97 |
| 5 | 15.55 | 10.58 | 10.45 |
| 10 | 7.22 | 5.67 | 4.50 |
| 15 | 3.38 | 2.53 | 2.12 |
| 20 | 1.18 | 1.31 | 0.64 |
| 25 | 0.64 | 0.91 | 0.29 |
| Number of pooled cells | **1093** | **1975** | **2019** |

**Table 8.** Frequency distribution of the distance between Pyy^+^ cells and Pgp9.5^+^ sensory neurons in the various parts of the colon in control and DSS-treated mice.

|  | Proximal colon | | Mid colon | | Distal colon | |
| --- | --- | --- | --- | --- | --- | --- |
| Distance (μm) | **Control** | **DSS** | **Control** | **DSS** | **Control** | **DSS** |
| 0 | 64.69 | 53.44 | 72.57 | 49.56 | 79.82 | 57.99 |
| 5 | 18.69 | 22.42 | 11.60 | 15.00 | 10.64 | 10.55 |
| 10 | 9.22 | 10.82 | 6.37 | 10.95 | 4.98 | 7.42 |
| 15 | 4.52 | 5.16 | 3.98 | 5.92 | 2.66 | 6.98 |
| 20 | 1.64 | 2.84 | 2.57 | 4.13 | 1.18 | 4.94 |
| 25 | 0.91 | 1.85 | 1.36 | 3.30 | 0.45 | 3.95 |
| 30 | 0.12 | 1.13 | 0.72 | 2.53 | 0.21 | 2.74 |
| 35 | 0.122 | 0.92 | 0.64 | 2.26 | 0.026 | 2.14 |
| 40 | 0.061 | 0.65 | 0.075 | 1.99 | 0 | 1.37 |
| 45 | 0 | 0.37 | 0.0758 | 1.84 | 0 | 1.04 |
| 50 | 0 | 0.27 | 0 | 1.22 | 0 | 0.54 |
| 55 | 0 | 0.063 | 0 | 1.25 | 0 | 0.27 |
| Number of pooled cells | **1637** | **4741** | **2636** | **3359** | **3712** | **1819** |

**Table 9.** Frequency distribution of the distance between Pyy^+^ cells and CD3^+^ T cells in the various parts of the colon in control and DSS-treated mice.

|  | Proximal colon | | Mid colon | | Distal colon | |
| --- | --- | --- | --- | --- | --- | --- |
| Distance (μm) | **Control** | **DSS** | **Control** | **DSS** | **Control** | **DSS** |
| 0 | 32.73 | 52.94 | 9.82 | 41.42 | 11.60 | 51.37 |
| 5 | 22.95 | 22.39 | 17.85 | 19.00 | 15.22 | 19.19 |
| 10 | 17.53 | 12.83 | 16.80 | 14.17 | 17.064 | 14.07 |
| 15 | 11.90 | 6.27 | 19.23 | 9.78 | 16.99 | 8.16 |
| 20 | 7.86 | 3.73 | 15.99 | 6.74 | 13.17 | 2.68 |
| 25 | 3.08 | 1.07 | 9.00 | 3.52 | 8.80 | 1.95 |
| 30 | 0.95 | 0.33 | 5.19 | 2.22 | 6.48 | 1.34 |
| 35 | 1.38 | 0.22 | 2.43 | 1.36 | 4.84 | 0.48 |
| 40 | 0.85 | 0.11 | 1.54 | 0.92 | 3.071 | 0.30 |
| 45 | 0.53 | 0 | 1.37 | 0.55 | 1.63 | 0.36 |
| 50 | 0.21 | 0.056 | 0.73 | 0.24 | 1.09 | 0.06 |
| Number of pooled cells | **941** | **1768** | **1252** | **1619** | **1465** | **1641** |
