## Supplemental Figures for "L-Cells are the Functional Neuropod Cell in Human Gastrointestinal Tract and are Dysregulated in Inflammatory Bowel Disease (IBD)"

### Slide 1
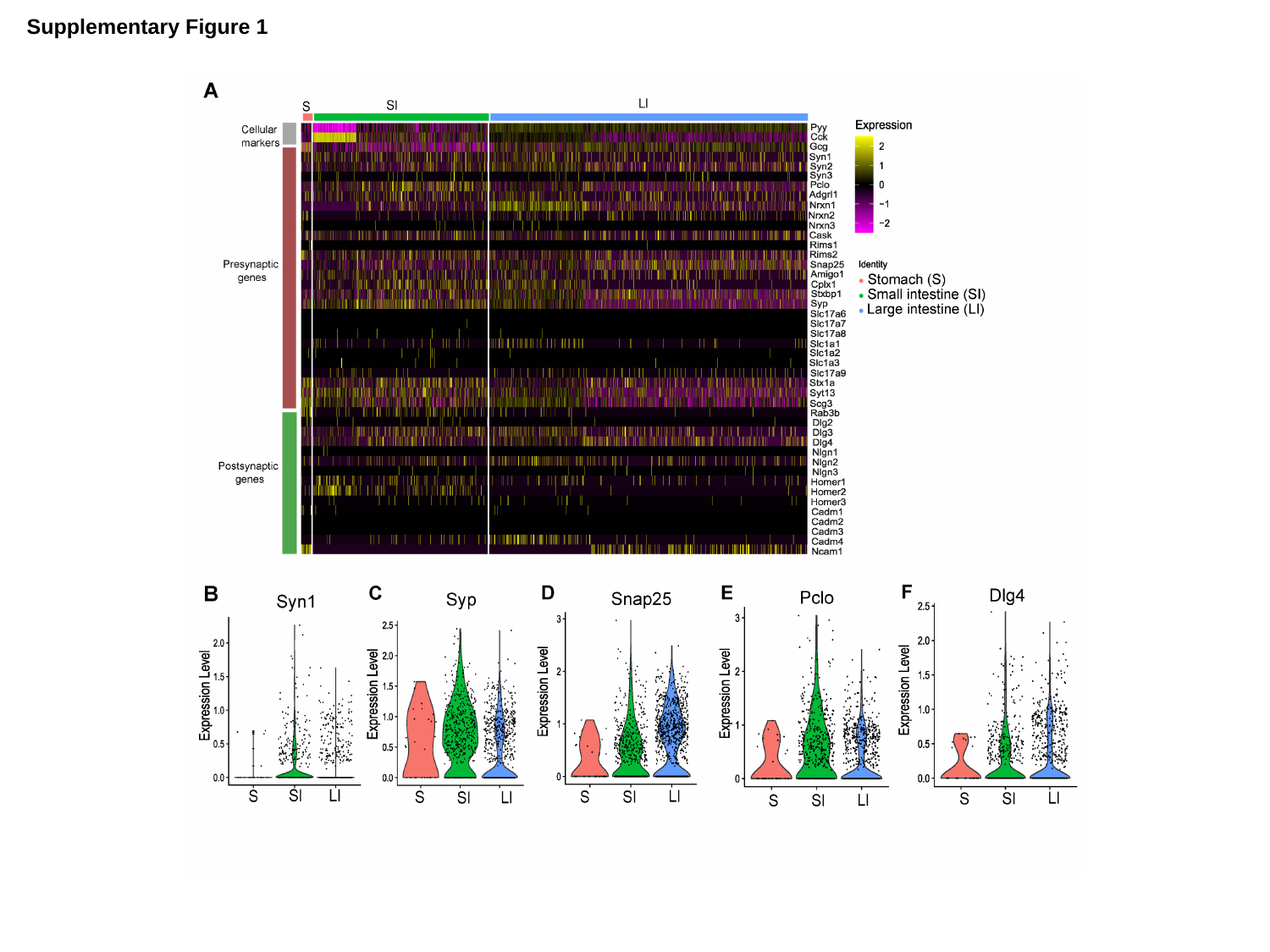

Supplementary Figure 1

### Slide 2
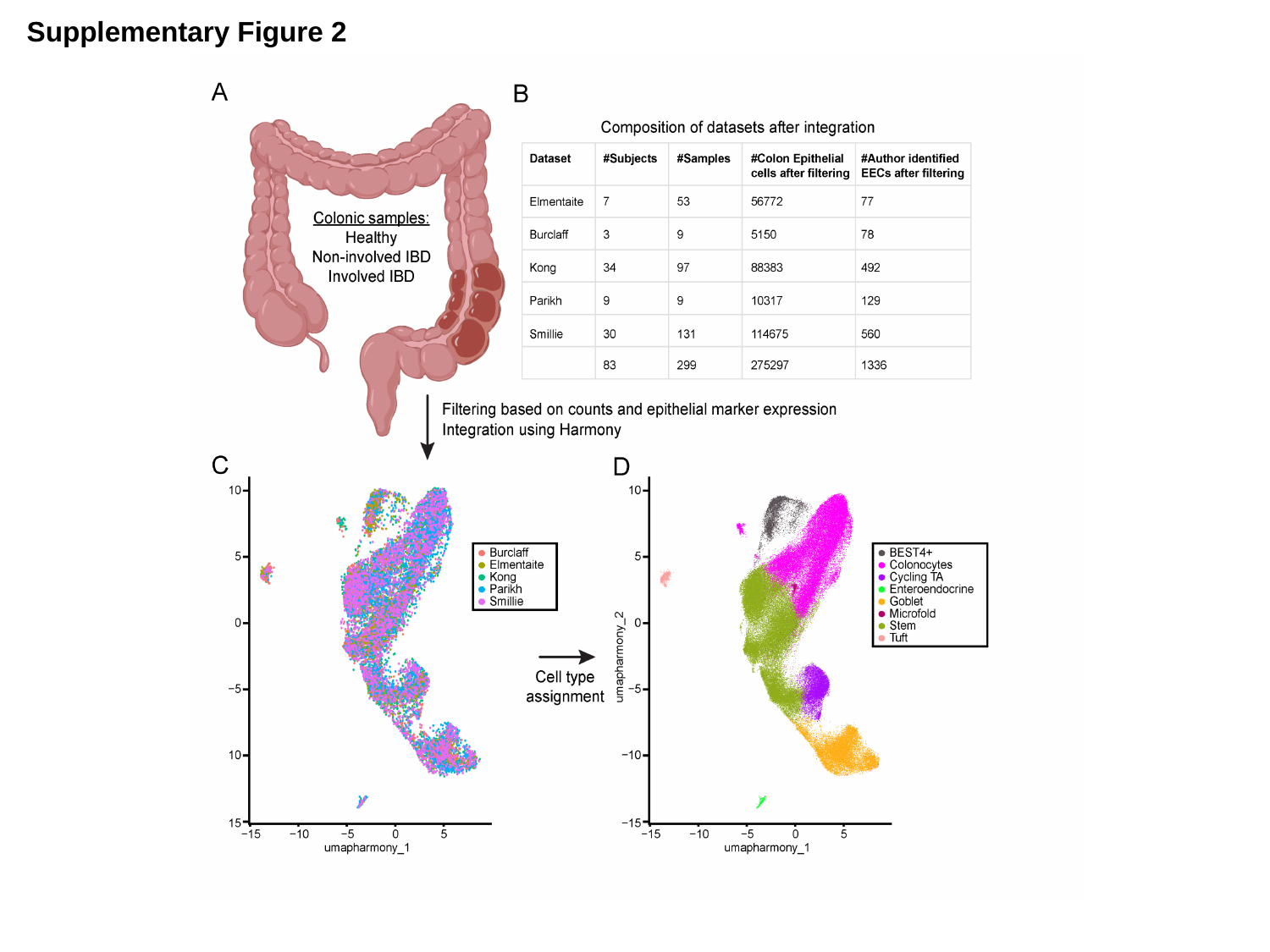

Supplementary Figure 2

### Slide 3
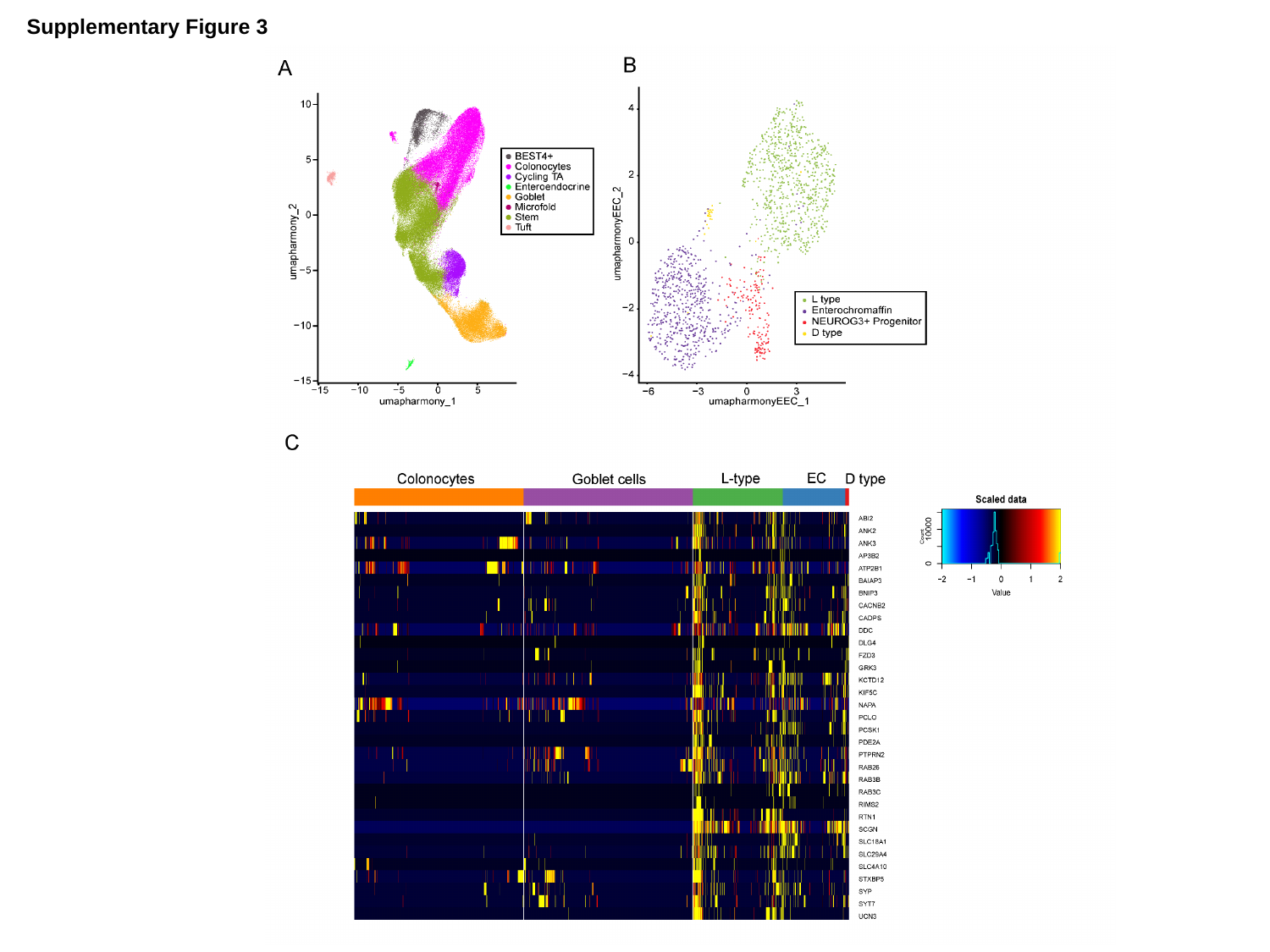

Supplementary Figure 3

### Slide 4
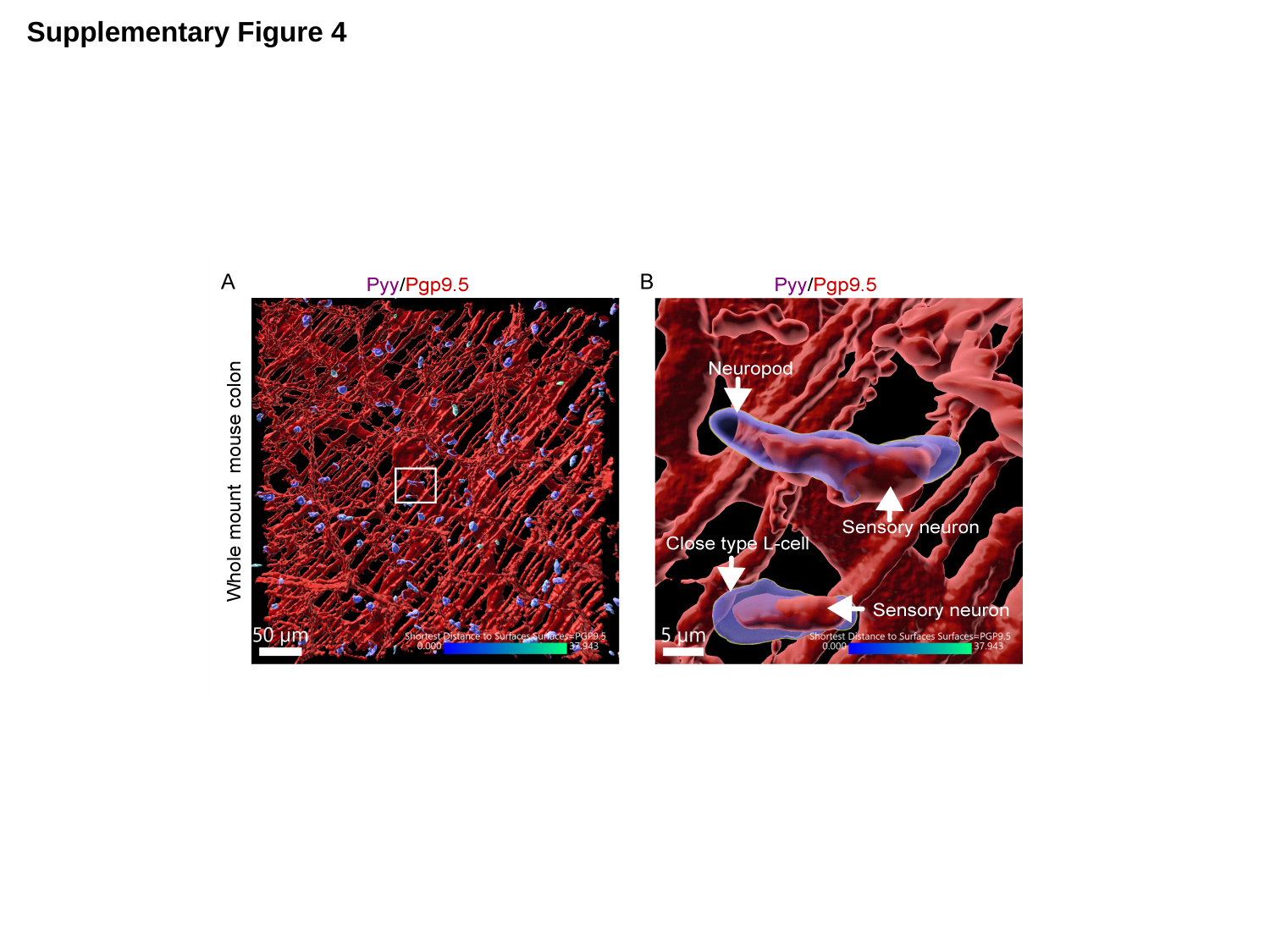

Supplementary Figure 4

### Slide 5
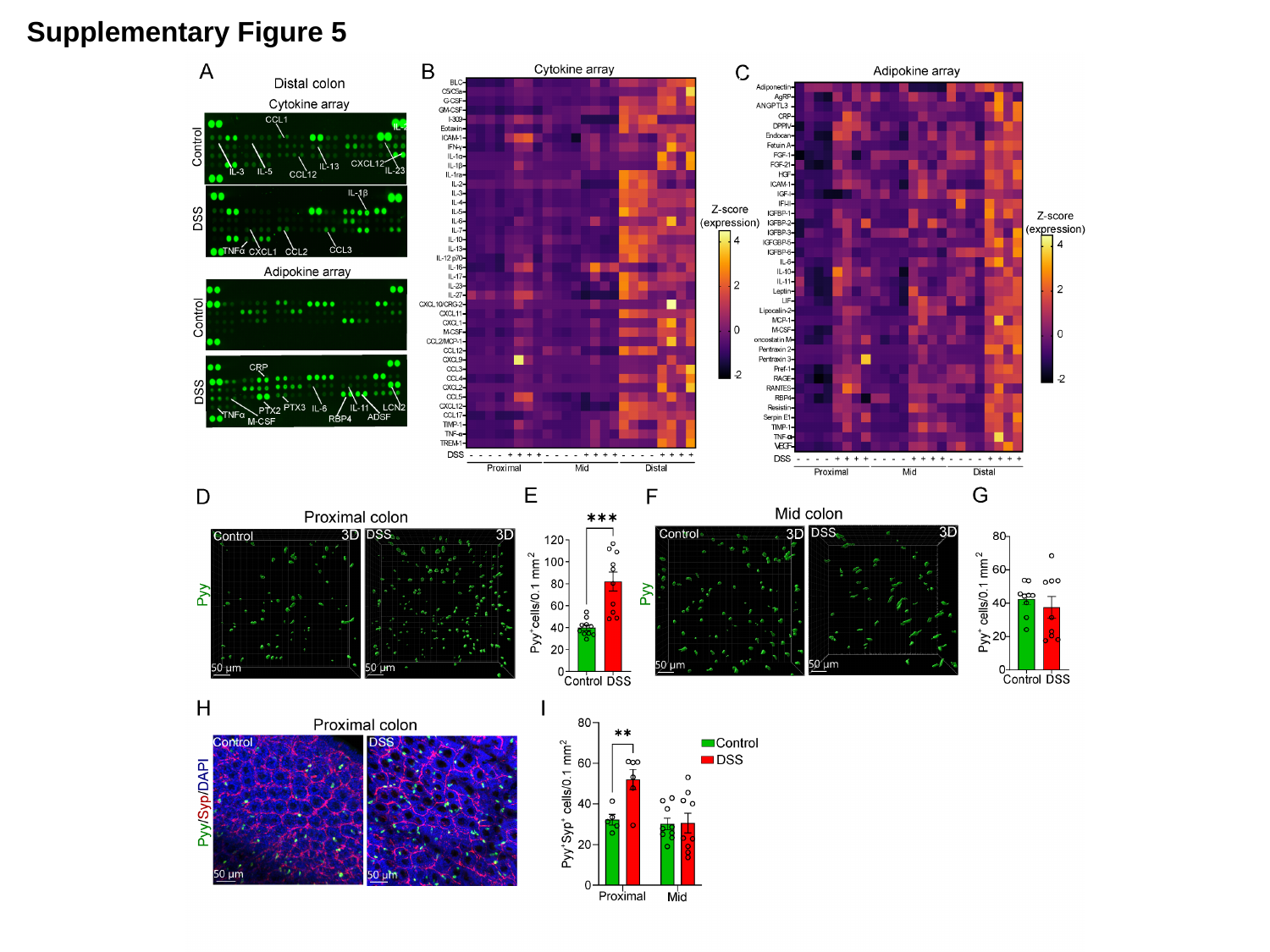

Supplementary Figure 5

### Slide 6
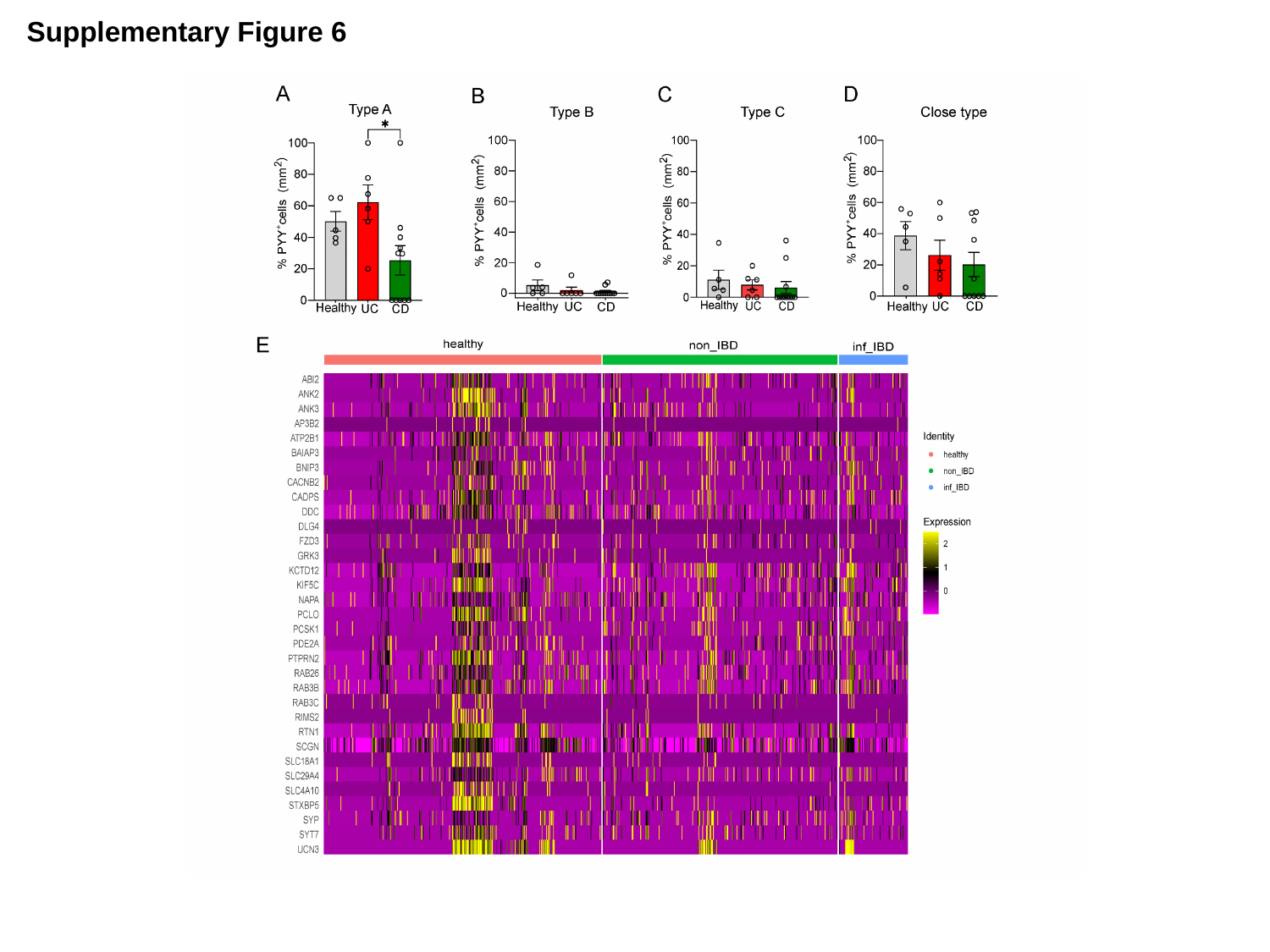

Supplementary Figure 6

### Slide 7
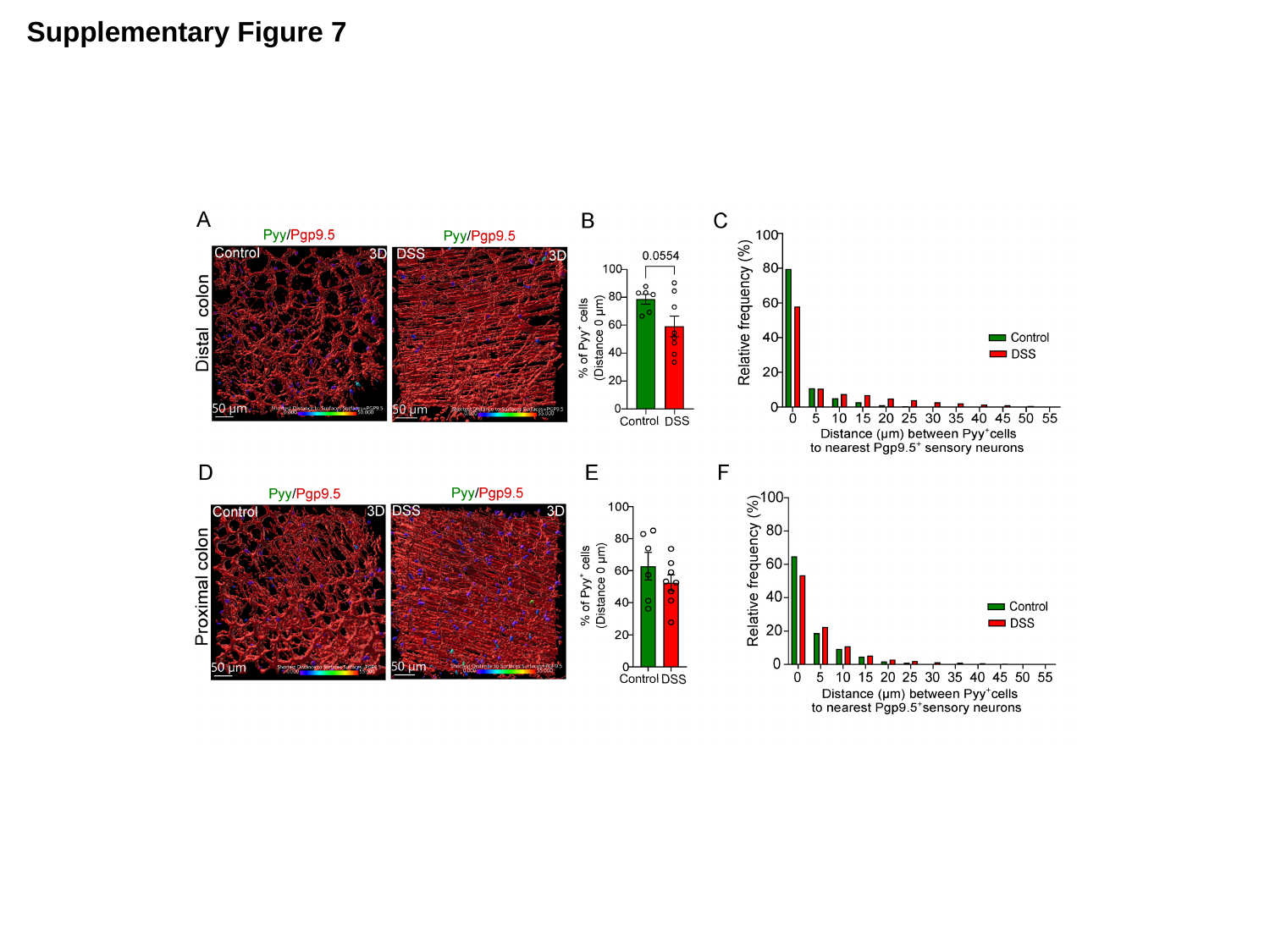

Supplementary Figure 7

### Slide 8
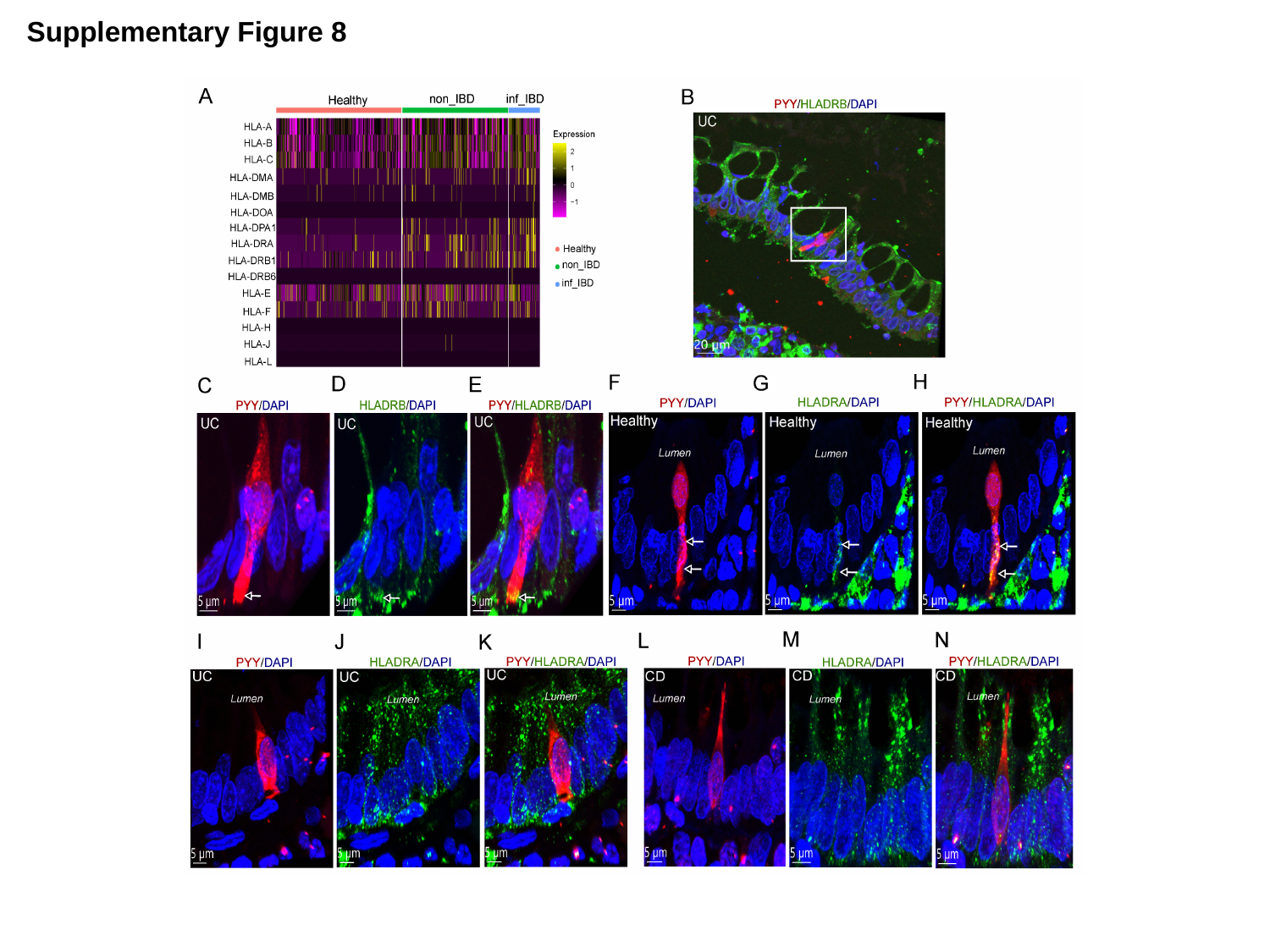

Supplementary Figure 8

### Slide 9
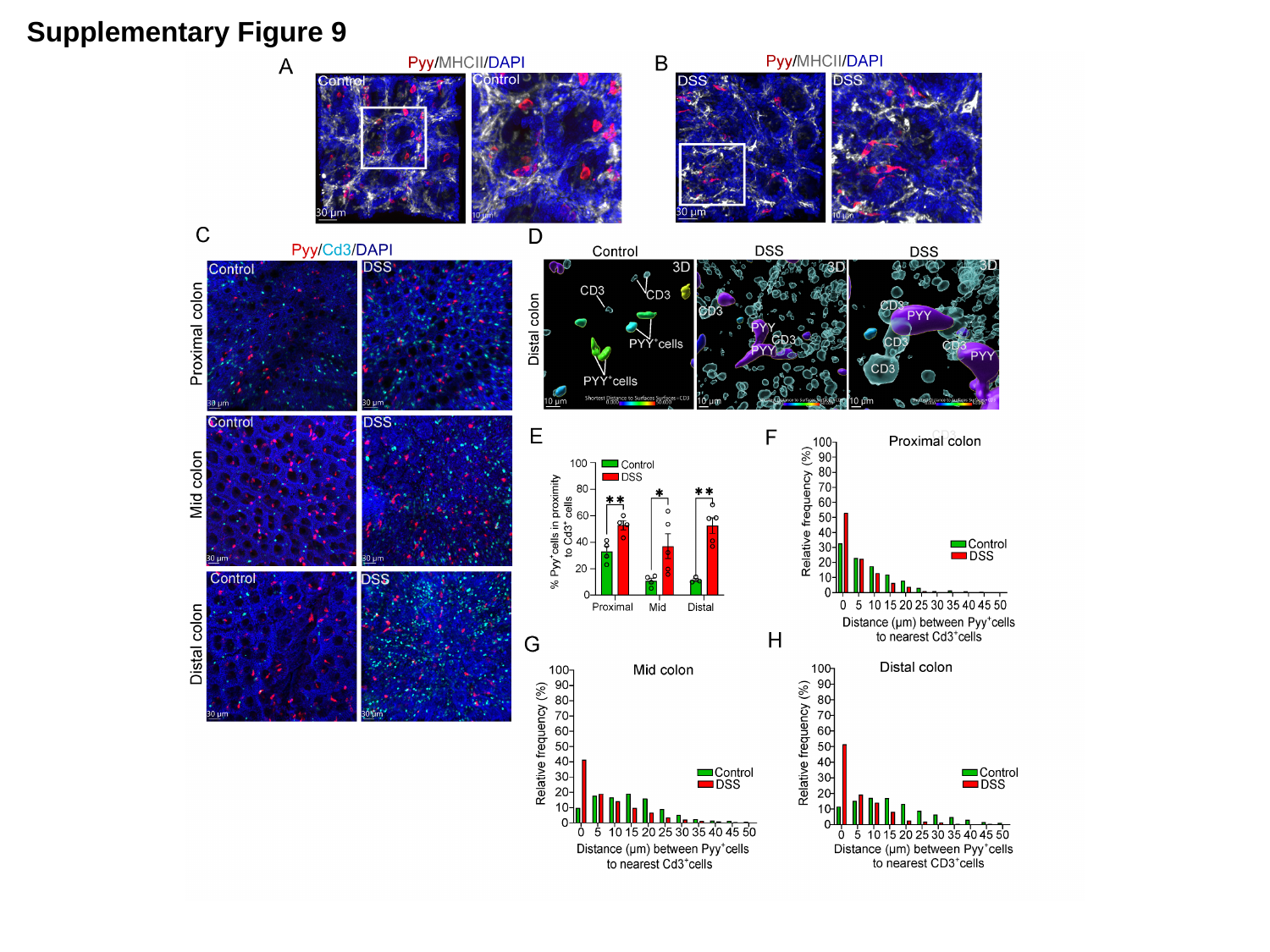

Supplementary Figure 9
